# Spatiotemporal patterns of neuronal subtype genesis suggest hierarchical development of retinal diversity

**DOI:** 10.1101/2021.04.29.442012

**Authors:** Emma R. West, Sylvain W. Lapan, ChangHee Lee, Kathrin M. Kajderowicz, Xihao Li, Connie L. Cepko

## Abstract

How do neuronal subtypes emerge during development? Recent molecular studies have expanded our knowledge of existing neuronal diversity. However, the genesis of neuronal subtypes remains elusive and previous studies have been limited by a lack of quantitative methods for simultaneous detection of subtype diversity *in situ*. The bipolar interneurons of the mammalian retina represent a diverse neuronal class, characterized by distinct functions, morphologies, and recently discovered transcriptional profiles. Here, we developed a comprehensive spatiotemporal map of bipolar subtype genesis in the retina. Combining multiplexed detection of 16 RNA markers with timed delivery of EdU and BrdU, we analyzed more than 30,000 single cells in full retinal sections to classify all bipolar subtypes and their birthdates. We found that bipolar subtype birthdates are ordered and follow a centrifugal developmental axis. Spatial analysis revealed a striking oscillatory wave pattern of bipolar subtype birthdates, and lineage analyses suggest clonal restriction on homotypic subtype production. These results inspired a hierarchical model of neuronal subtype genesis in the mammalian retina, with the wave pattern of subtype birthdates arising from early asymmetric cell divisions among founding retinal progenitor cells. Our results provide an outline of the developmental logic that generates diverse neuronal subtypes, and establishes a framework for studying subtype diversification.

## 1. Introduction

Recent technological advancements have enabled the systematic discovery of cellular diversity at unprecedented resolution. Importantly, single-cell RNA sequencing has provided quantitative definitions of cellular diversity by uncovering unique and measurable transcriptional profiles. Retinal neurons are an ideal model for studying neuronal diversification, as they display modular neuronal subtype organization with well-defined transcriptional and morphological distinctions. Six major neuronal cell types are present in the mammalian retina, with extensive subtypes for specialized visual circuitry. Recent work has identified >100 neuronal subtypes in the mouse retina, many of which are conserved in primates and humans [1–4].

Among the diverse retina neurons, murine bipolar interneurons are classified into 15 subtypes with characteristic morphologies, transcriptomes, and functions. Bipolar interneurons bridge all visual circuits, connecting sensory rod and cone photoreceptors to the output neurons, retinal ganglion cells (Fig. 1A). They are the latest-born neuronal cell type, exiting mitosis during the first postnatal week after most other retinal neurons have already differentiated, and they do not migrate from their position of origin [5]. Despite extensive molecular profiling of bipolar subtypes, the development of bipolar cell diversity is not understood. To date, no studies have simultaneously examined all bipolar subtypes in the same tissue, due to a lack of quantitative methods for subtype identification and the technical limitations of combinatorial marker detection.

**Figure 1.**
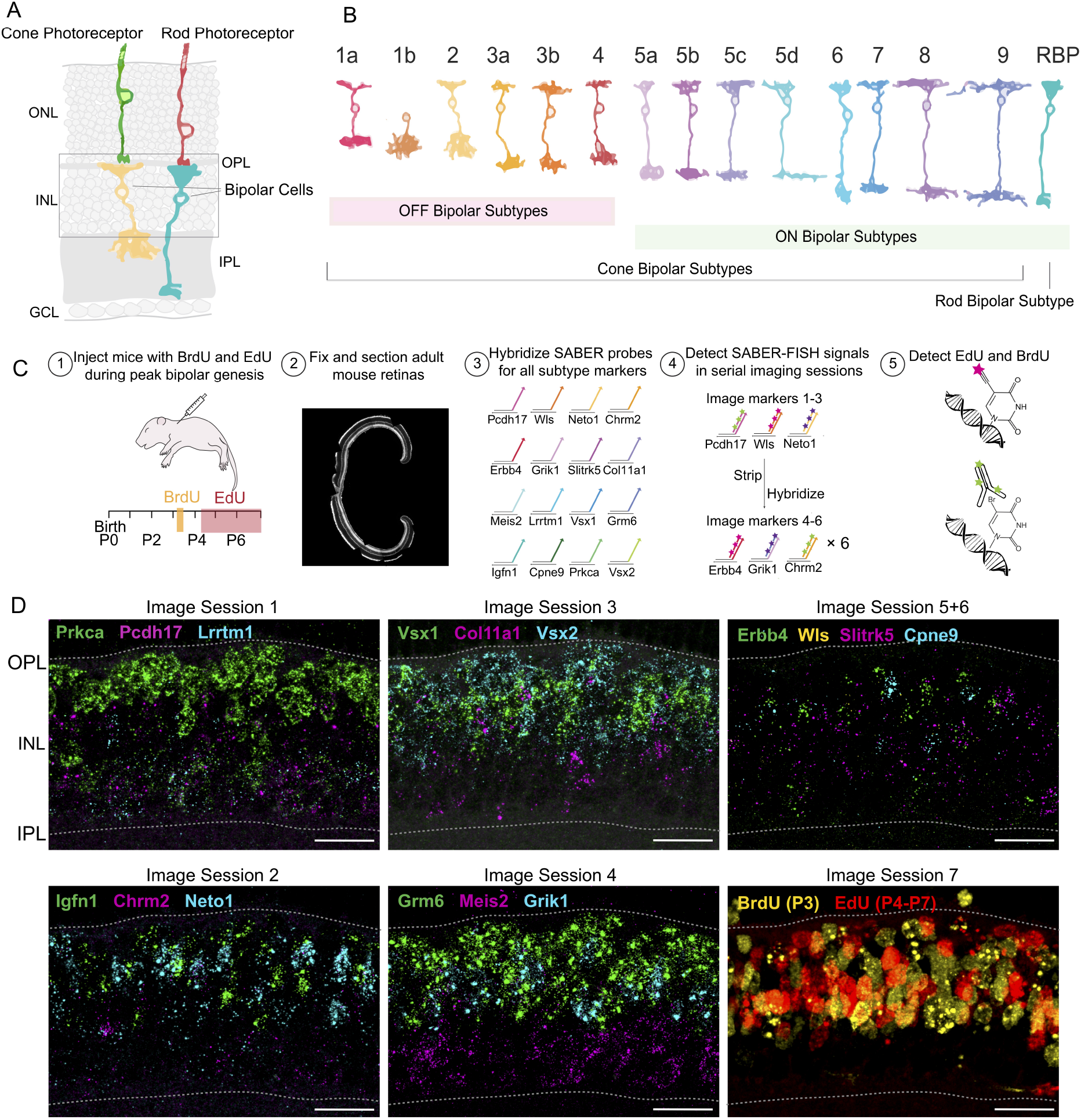
Multiplexed SABER-FISH combined with EdU and BrdU detection in tissue sections labels all bipolar subtypes and their birthdates. **(A)** Schematic of a transverse retina section with bipolar cells connecting to rod and cone photoreceptors. Bipolar cell bodies are located in the inner nuclear layer (INL), with their dendrites in the outer plexiform layer (OPL) forming synapses with photoreceptors. Different bipolar subtypes laminate within different sublaminae of the inner plexiform layer (IPL), contacting different visual circuitry. **(B)** Figure adapted from [4]. Fifteen subtypes of bipolar cells have been classified based on their transcriptional profiles, morphologies, and function. The OFF subtypes (1a-b, 2, 3a-b, 4) hyperpolarize in response to light, while ON subtypes (5a-d, 6, 7, 8, 9, RBP) depolarize to the same stimuli. There are 14 types of cone bipolar cells and 1 type of rod bipolar cell. **(C)** (1) Mice were injected with EdU and BrdU during bipolar cell genesis, with a pulse of BrdU on P3, and EdU pulses every 12 hours through P7. (2) P18 eyes were dissected, fixed, frozen, and cryosectioned along the DV axis. A representative section is shown, stained with WGA. (3) Sixteen marker genes were chosen which display differential expression among single cell Drop-Seq profiles for the 15 bipolar subtypes [4]. Each marker gene was targeted by SABER-FISH probes with a unique concatamer sequence, allowing each marker to be independently detected and imaged. (4) SABER-FISH signal for each marker RNA was detected in 6 serial image sessions, with up to 3 markers imaged per session. For each session, short fluorescent oligonucleotides were hybridized to the marker-specific SABER probes. After imaging, these oligonucleotides were removed by a stringent wash, and fluorescent oligos targeting the next subset of markers were hybridized. (5) After SABER-FISH imaging, EdU and BrdU were detected and imaged. **(D)** Representative maximum intensity projections of SABER-FISH data for each bipolar subtype marker gene and BrdU and EdU. Scale bars are 25μm

In this study, we aimed to describe the patterns of bipolar genesis of all subtypes at high spatiotemporal resolution by leveraging published molecular profiles for subtype identification. Using a combination of multiplexed RNA FISH and dual-nucleoside analog ‘window labeling’ to mark times of mitotic exit, we investigated the relationship between bipolar subtype identity, location, and birthdate for all known subtypes in the same retinas. We classified all 15 bipolar subtypes by co-detecting 16 RNA markers and employed automated 3-D image analysis of more than 20,000 bipolar cells to discover continuous patterns of subtype genesis in entire retina sections.

Our results demonstrate that bipolar subtype birthdates are distinct but overlapping, and confirm that their birth order correlates with their role in visual function. We found that bipolar birthdates follow a central-to-peripheral pattern that progresses independently of the central retina, and that subtype birthdates arrange in a wave pattern across the retina, with local subtype genesis oscillating in space and time. Lineage analysis of postnatal clones showed a bias against homotypic subtype composition, providing evidence for fine-tuned regulation of local subtype genesis. Together, these results impel a hierarchical model of bipolar subtype production with the oscillatory wave pattern arising as a result of early asymmetric divisions among founding retinal progenitor cells.

## 2. Results

### 2.1. Capturing the complete diversity of bipolar subtypes in situ

Previous studies of bipolar cells have been limited to only a handful of subtypes, due to a lack of subtype-specific marker genes and the limited ability to co-detect those markers *in situ*. Recently, single cell RNA sequencing of mature mouse bipolar cells uncovered transcriptional signatures for all bipolar subtypes, 15 in total, providing markers to identify each subtype (Fig. 1B). Leveraging this, we aimed to simultaneously classify all bipolar subtypes in tissue sections to capture the spatiotemporal patterns of their genesis at cellular resolution.

We combined BrdU and EdU labelling of cellular birthdates with multiplexed detection of bipolar subtype markers in situ (Fig. 1C). To visualize bipolar subtypes, we detected 16 RNA markers with validated differential expression across the 15 subtypes using serial multiplexed SABER-FISH in retinal tissue sections [6] (see Methods). To label bipolar cell birthdates, mice were injected with a pulse of BrdU at postnatal day 3 (P3), followed by a series of EdU injections through the duration of bipolar genesis (P4-P7). Retinas were harvested at P18, when bipolar subtypes are terminally differentiated, for detection of the incorporated nucleoside analogs and subtype marker RNAs (Fig. 1D). Successful co-detection of BrdU and EdU with subtype markers *in situ* permits overlay of cellular birthdates, subtype identities, and tissue location.

With the goal of analyzing thousands of bipolar cells across large retina regions, we developed a pipeline to automatically classify bipolar subtypes in situ (see Methods) (Fig. 2A, Supplementary Fig. 1A). SABER-FISH image data were converted to single-cell gene expression profiles of imaged cells. Using these data from 14,534 imaged cells (Fig. 2B-D, Supplementary Fig. 2B-C), we trained a random forest classifier to identify bipolar subtypes in 3-D tissue sections. When tested on 1,000 manually classified cells, the subtype classifier performed with an F1 score of 0.93, confirming that the automated classifications closely matched manual classifications (Fig. 2E). The classified bipolar subtype clusters had normalized gene expression profiles consistent with previous single cell data with Pearson correlation coefficients of 0.9273 (*p* = 1.1046 × 10^-96^), 0.9411 (*p* = 1.5431 × 10^-106^), and 0.9470 (*p* = 1.9064 × 10^-111^) for three retina replicates (Fig. 2F, Supplemerary Fig. 1D-E). Close inspection of individual classified cells confirmed that marker expression patterns of each subtype cluster reflect the expression patterns in single cells (Fig. 2G). Type 9s are known to express Grm6, Cpne9, and Vsx2, while not expressing Grik1, Neto1, and Prkca, for example (Fig. 2G, first row). Similarly, Type 2s are known to express Neto1, low levels of Grik1, Vsx1, and Vsx2, but not Igfn1 or Cpne9 (Fig. 2G, bottom row). Despite fundamental differences in methodology for RNA detection, the relative RNA expression patterns across bipolar subtypes were consistent between SABER-FISH profiles and published single-cell expression profiles.

**Figure 2.**
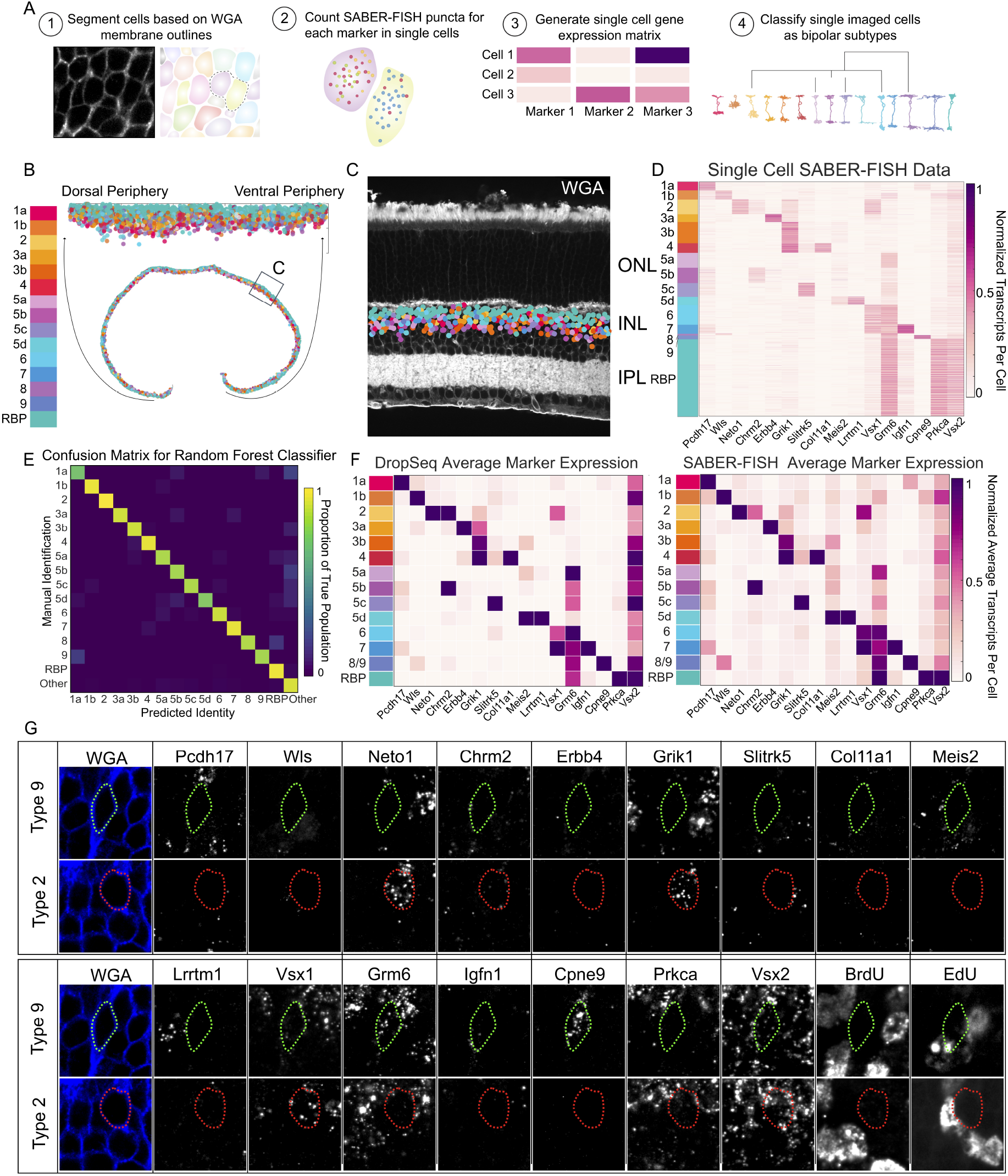
Automated classification of all bipolar subtypes *in situ*. (**A**) Computational pipeline for identifying bipolar subtypes *in situ* based on SABER-FISH RNA marker expression data. (1) Cells were segmented in 3-D based on a wheat germ agglutinin (WGA) membrane stain. (2) SABER-FISH fluorescent puncta for each marker gene were counted within each cell. (3) SABER-FISH puncta per cell counts and integrated fluorescent intensity per cell were used to construct a single cell gene expression matrix. (4) Gene expression data were used to train a classifier to identify bipolar subtypes. (**B**) Example retina section with classified bipolar subtypes along the DV axis shown as dots colored by subtype. (**C**) Example region of the retina with classified bipolar cells shown as dots colored according to their subtype identity. (**D**) Heatmap of normalized SABER-FISH puncta per cell for each marker gene across subtype clusters. These data were used to train a subtype classifier. Values were normalized by the maximum for each gene across all bipolar subtype clusters. (**E**)Heatmap of average SABER-FISH puncta detected for each marker gene across bipolar subtype clusters (see Methods). For plotting, values were normalized by the maximum for each gene across all bipolar subtype clusters. (**F**) Heatmap of average Drop-Seq transcripts per cell detected for each marker gene across bipolar subtype clusters. Values were normalized by the maximum for each gene across all subtype clusters. (**G**) Examples of two single cells classified as Type 9 (green outline) and Type 2 (red outline) bipolar cells. SABER-FISH images for all subtype markers are shown for a single plane.

Bipolar subtype abundances vary widely, with a 10 to 15-fold difference in cellular density between the most abundant (RBPs) and rarest (Type 9) subtypes [7]. The captured subtype densities were consistent with those found in serial electron microscopy reconstructions [7] (Supplementary Fig. 1F). We also asked whether the soma location of our classified subtypes were consistent with known patterns within the inner nuclear layer (INL) of the retina. Consistent with previous reports, we observed the rod bipolar population located more proximal to the outer plexiform layer than cone bipolar subtypes (Supplementary Fig. 1G).

Local bipolar cell densities were calculated to validate spatial reconstruction across the dorsoventral (DV) axis. The average density of classified bipolar cells was 40,592 ± 5,806 bipolar cells/mm^2^ (mean ± standard deviation across retinas), which agrees with the previous estimate of 39,975 ± 6,339 bipolar cells/mm^2^ based on a combination of transgenic mouse lines and type-specific antibodies (Supplementary Fig. 1H) [8].

Slightly higher density of OFF subtypes were observed in the ventral retina compared to the dorsal retina, consistent with previous analyses (Supplementary Fig 1I) [9]. Similarly, slightly higher ventral density of Type 9 was observed, consistent with a recent observation of ventral enrichment for this S-cone selective subtype (Supplementary Fig. J-K) [10]. Overall, the classified bipolar subtypes matched published gene expression and anatomical data, permitting analyses of the full diversity of bipolar cells in situ at scale.

### 2.2. Bipolar subtypes are born at overlapping but distinct times

The major classes of retinal neurons are born in distinct, overlapping windows of genesis, in a conserved order across species (reviewed in [11]). Few studies have examined birthdate patterns of retinal neuron subtypes, and none have captured patterns for all subtypes within a neuronal class. One study established that cone bipolar genesis commences before rod bipolar genesis but that their periods of genesis significantly overlap [12,13]. Similarly, coarsely separated amacrine subtypes have distinct but overlapping genesis windows. We set out to discover whether individual bipolar subtypes have different birthdates and whether they are born in a pattern that might shed light on how bipolar diversity is established.

Previous studies have shown that bipolar cells are born during the first postnatal week, with peak genesis around P3 [14]. To label bipolar cell birthdates (the time of terminal mitosis), we used a strategy to label dividing cells with nucleoside analogs. We first injected BrdU on P3 to label cells in S-phase at the peak of bipolar birthdates. We then injected EdU every 12 hours through the first postnatal week, starting 16 hours after the initial BrdU injection. The timing of EdU injections was chosen based on known cell cycle kinetics [14] to label all cells born after the first EdU injection on P4 (Supplemetary Fig. 2A). This strategy allows us to bin all cells into three populations: cells born before P3 are BrdU^-^/EdU^-^, cells born in the 16 hour window between P3 and P4 are BrdU^+^ /EdU^-^, and cells born after P4 are either BrdU^+^ /EdU^+^ or BrdU^-^ /EdU^+^ (Fig. 3A-B, Supplemetary Fig. 2B). Using this precisely timed series of BrdU and EdU injections and automated automated segmentation to quantify the BrdU and EdU signal in single cells (Supplemetary Fig. 2C), birthdate information was captured about every imaged cell, with temporal precision of 16 hours during peak bipolar genesis.

**Figure 3.**
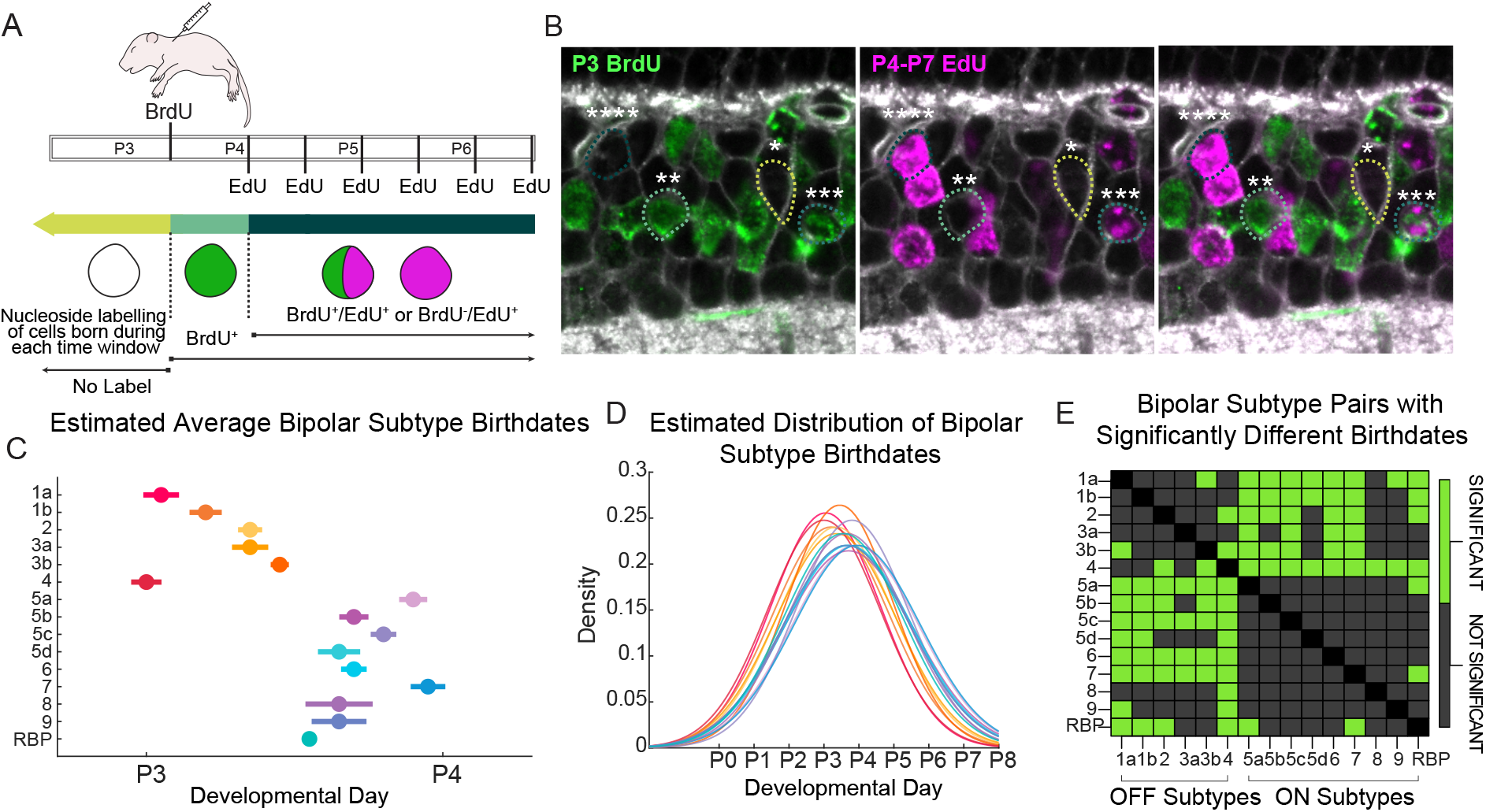
Bipolar subtypes have distinct but overlapping birthdates. (**A**) Pups were injected with BrdU on P3, followed by EdU 16 hours later and every 12 hours through the end of retinal proliferation on P7. Retinas were harvested at P18 and stained for subtype markers and nucleoside analog labels. Colors correspond to binned windows of terminal S-phase completion, i.e. birthdate. All cells fall in one of three bins: those that completed terminal S-phase before BrdU injection (yellow, BrdU^-^ /EdU^-^), those that completed terminal S-phase between BrdU injection and the first EdU injection (green, BrdU^+^ /EdU^-^), and those that completed terminal S-phase after the first EdU injection (blue, [BrdU^+^ /EdU^+^; BrdU^-^ /EdU^+^]). (**B**) Example of BrdU and EdU detection in retina section. Cells with each of the four possible nucleoside labels are outlined: (*) = BrdU^-^ /EdU^-^, (**) = BrdU^+^ /EdU^-^, (***) = BrdU^+^ /EdU^+^, (****) = BrdU^-^/EdU^+^. (**C**) Estimated average birthdates for each subtype. Error bars reflect the estimated standard error of the mean. (**D**) Estimated probability density function predicting the distribution of birthdate for each subtype during the first postnatal week (see Methods). (**E**) Binomial z-tests were performed to compare the proportion of each pair of bipolar subtypes born before P4. For each subtype, the fraction of cells born before P4 across all retinas was calculated by summing the number of EdU^-^ /BrdU^-^ cells + BrdU^+^ /EdU^-^ cells, and dividing by the total number of cells with that subtype identity. Matrix shows p-value results for each tested pair of subtypes, with a threshold of 0.05 for significance after Bonferroni correction for multiple hypothesis testing. (green = birthdates are significantly different between the subtype pair, grey = no significant difference was observed).

Since the raw birthdate data are discreet with respect to time, a model was fit to estimate the continuous birthdate patterns for each subtype over developmental time (see Methods). The model estimates the average birthdate for each subtype and the standard deviation of birthdate, deriving a probability density function (PDF) for subtype production on each day between P0 and P8. The estimated average birthdate for each subtype falls between P3 and P4 but is ordered within this range (Fig. 3C). Overlaying the estimated PDFs for each subtype illustrates that their genesis is highly overlapping with all subtypes born during the first postnatal week (Fig. 3D).

Despite the large overlap in bipolar subtype genesis, direct comparison between subtypes revealed that many have significantly different birthdate patterns (Fig. 3E-F). For example, more than 60% of Type 4s were born before P4, while only 40% of Type 7s were born before P4. For pairwise comparison of subtype birthdates, binomial z-tests confirmed that many of the observed differences were statistically significant (Fig. 3E). Grouping all OFF and ON bipolar cells, 57.6% of OFF bipolar cells were born before P4, compared to 48.6% of ON bipolar cells (*p* = 2.2 × 10^-16^). In general, individual OFF subtypes were born significantly earlier than the ON subtypes, and Types 4 and 1a have the earliest birthdates (Fig 3E-F). Birthdates of at least four cone bipolar types (1a, 1b, 2, and 4) significantly precede RBP birthdates, consistent with previous data [12] (Fig. 3F).

Some subtype pairs, however, did not show significantly different birthdates. For example, we were unable to statistically resolve the birthdates of Types 5a-d or Types 2 and 3a (Fig. 3F). This could reflect either that these types are truly born at the same time or that their genesis is extremely overlapping, requiring more statistical power to resolve such fine differences.

Since it is known that bipolar subtypes are present at very different numbers in the mouse retina [8], we asked whether more prevalent subtypes were born over longer periods of time, or whether more cells are produced during a comparable time. The estimated standard deviation in birthdate for each subtype did not correlate with subtype prevalence (*ρ*=-0.0713, *p*=0.8005), suggesting that the production of more numerous cells is not achieved by significantly extending the genesis window, but by making more cells in the same span of time.

Taken together, these results indicate that bipolar birthdates are ordered for individual subtypes, and that OFF bipolar subtypes are generally earlier-born than ON subtypes. At any given time, however, many different subtypes can be concurrently born, necessitating mechanisms for choosing among the diverse bipolar subtype fates at a given space and time.

### 2.3. Bipolar birthdates follow a central-to-peripheral genesis pattern that does not require the central retina

Previous studies have demonstrated that retinogenesis proceeds in a central-to-peripheral gradient, with developmental events occurring in the central retina before the periphery. In rodents, the birth of retinal neurons begins centrally and spreads peripherally with a delay of 2-3 days [14,15]. It has been shown that some subsets of amacrine cells display the same centrifugal genesis pattern [16], but the pattern of bipolar subtype genesis has not been systematically described for all subtypes.

To assess whether individual bipolar subtypes follow a central-to-peripheral pattern, we compared the birthdates of central versus peripheral cells of each subtype. In all cases, the central bipolar population was born earlier than the peripheral population (Fig. 4A). On average, 63% of central bipolar cells were born before P3, in contrast to only 19% of peripheral bipolar cells. These data confirm that all bipolar subtypes are born in a centrifugal gradient.

**Figure 4.**
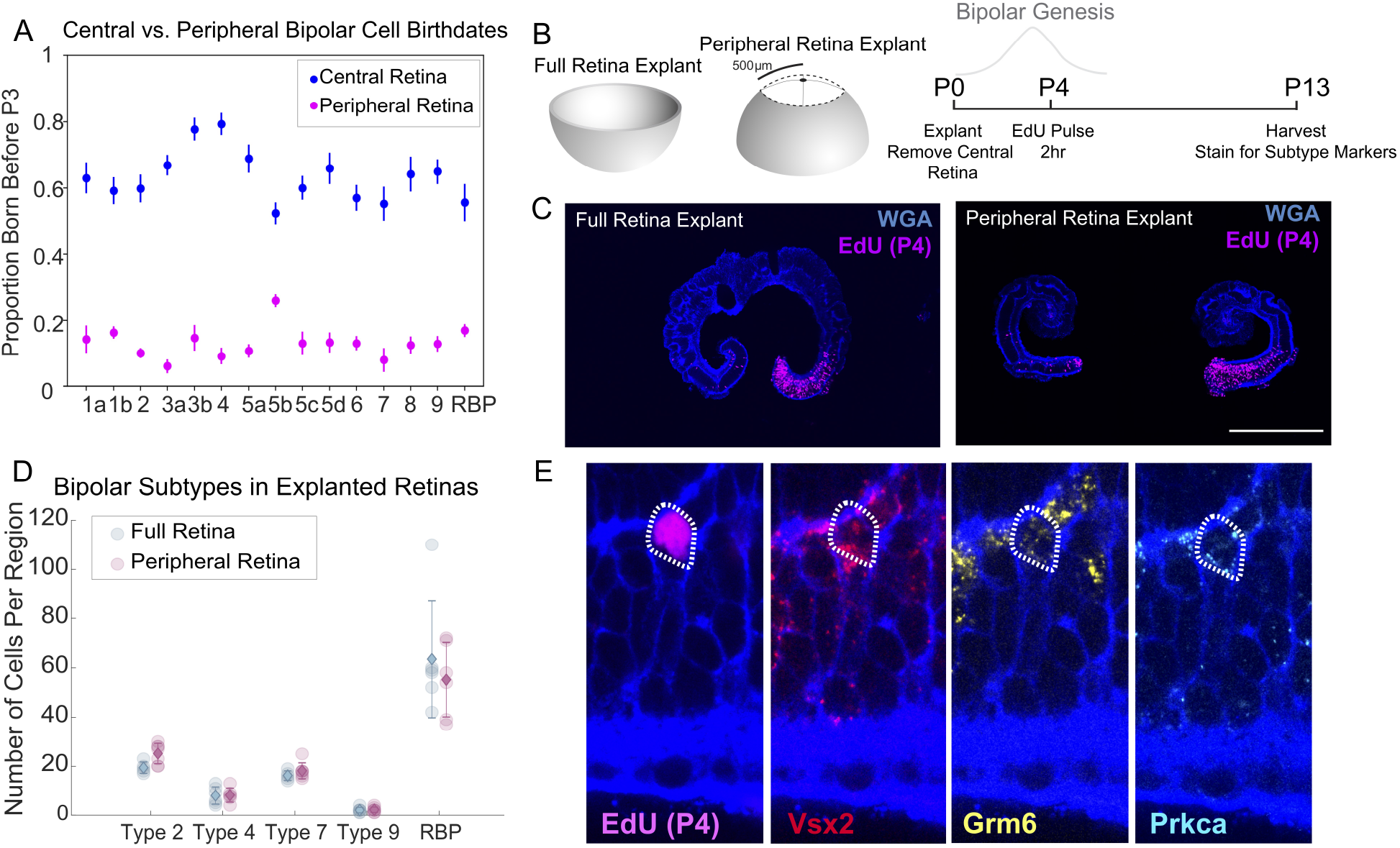
Bipolar subtypes are born in a central-to-peripheral pattern that progresses without the central retina. **(A)** Comparison of the average proportion of each subtype born before P3 in the central vs. peripheral retina. Error bars reflect standard error of the mean across three retina replicates. **(B)** Experimental schematic. Retinas were explanted from P0 mice and were cultured with or without removal of their center (1mm disk). On the 5th day of culture (P4), EdU was added to the media for 2 hours. Retinas were cultured for an additional 9 days before fixing and staining for SABER-FISH detection of bipolar subtype markers. **(C)** Example of sections from a full retina explant (top) and peripheral retina explant (bottom) at P13. Explants are stained for WGA (blue) and EdU (magenta). The central retina is positioned at the top of the image. Scale bars are 500μm. **(D)** SABER-FISH images for a subset of bipolar subtype markers in peripheral-only retina explant cultures. **(E)** Quantification of the number of cells with SABER-FISH expression profiles corresponding to Types 2, 4, 7, 9, and RBPs in explanted retinas. The number of each subtype was quantified in peripheral regions from whole retina explants (blue dots) and explants where the central retina was removed at P0 (purple dots). Diamonds represent the mean for each subtype, with error bars reflecting standard deviation. Six imaged regions from three different animals were analyzed for each condition. **(F)** Example of a RBP cell born in a peripheral retina explant after P4. SABER-FISH images for individual RBP marker genes are shown.

The central-to-peripheral birthdate pattern suggests that bipolar genesis in the peripheral retina might be induced or instructed by earlier born cells in the central retina. To test whether genesis and differentiation of bipolar subtypes in the peripheral retina depends on continuous cues from the central retina, we performed explant experiments where the central retina was removed at P0, days before most peripheral bipolars are born (Fig. 4B, show full in supplement). P0 retinal explants were cultured for two weeks either with or without removal of their center, and received a 2 hour EdU pulse at P4. After two weeks, 9 bipolar subtype markers were detected by SABER-FISH.

Explants with and without their centers continued to proliferate until at least P4, as evidenced by EdU incorporation (Fig. 4C). Normal subtype-specific gene expression patterns arose in the peripheral retina even when the center was removed (Fig. 4D). Cells in the periphery were present with subtype-specific marker expression patterns for Types 2, 4, 7, 9, and RBPs at indistinguishable frequencies with and without the central retina removed. These subtypes represent examples of OFF (Types 2 and 4), ON (Types 7, 9, and RBP), cone-connecting (Types 2, 4, 7, and 9), rod-connecting (RBPs), rare (Type 9), and prevalent subtypes (RBPs), all of which developed in the absence of the central retina. Importantly, EdU incorporation at P4 in some peripheral bipolar cells demonstrates that even cells born 5 days after removal of the central retina can undergo normal subtype differentiation (Fig. 4E).

### 2.4. Bipolar birthdates arrange in a wave-like pattern across the retina space

The injections of BrdU and EdU were timed based on known cell cycle kinetics [14], such that all cells in S phase at 17:00hr on P3 were labelled with BrdU, and all cells that underwent terminal S-phase after 09:00hr on P4 were labelled by EdU. Therefore, EdU-/BrdU- cells were born before P3, during the first half of bipolar genesis. BrdU+/EdU- cells were in S-phase at the time of P3 BrdU injection but exited mitosis after the following M-phase, and thus were born on P3-P4. BrdU+/EdU+ cells were similarly in S-phase on P3 during the BrdU label, but went on to divide at least one more time, incorporating EdU during a subsequent S-phase, and were in a non-terminal S-phase on P3-P4. Finally, BrdU-/EdU+ cells were not in S-phase at 17:00hr on P3, and were born after P4.

Subtype birthdate patterns were computed along the DV axis by calculating the local proportion of each subtype with each nucleoside analog label as a sliding window average across space. Comparing cells born before versus after P3, a striking wave pattern emerged of locally correlated birthdates of bipolar subtypes (Fig. 5B, Supplementary Fig. 3A-B). For all subtypes in all retinas, the DV axis was tiled by regularly spaced “peaks” of enrichment for early bipolar cell birthdates, adjacent to “troughs” with fewer early-born bipolar cells. The patterns were correlated among subtypes in the same retina, both globally and locally (Supplementary Fig. 3C), and overlay of the peaks called for different subtypes demonstrates roughly consistent peak locations (Fig. 5C, Supplementary Fig. 3B).

**Figure 5.**
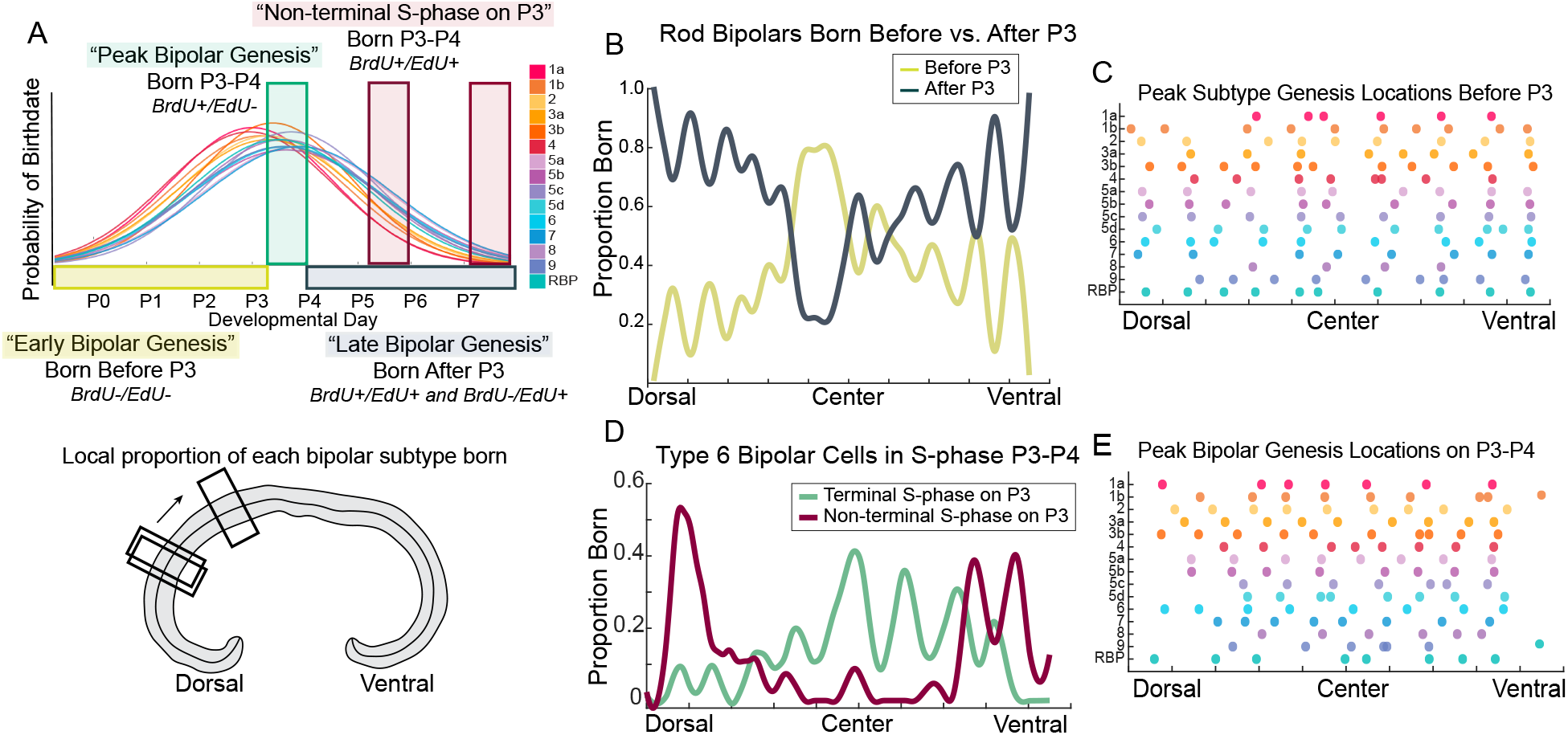
Bipolar subtype birthdates arrange in a wave pattern across the retinal dorsoventral axis. (**A**) (Top) Schematic of cellular birthdate labels based on timed BrdU and EdU injections, shown over the birthdate probability curves for each subtype during the first postnatal week (from Fig. 3D). Cells born before P3 are BrdU-/EdU- (yellow). Cells born after P4 are BrdU+/EdU+ or BrdU-/EdU+ (blue). Cells born on P3-P4 are BrdU+/EdU- (green). The BrdU+/EdU+ populations mark cells in S-phase on P3, which continued dividing after P4. (Bottom) The local proportion of each subtype born during each labelled time was computed across the DV axis as a sliding window average (SWA) (see Methods). (**B**) Loess-smoothed SWA of the local proportion of RBPs born across the DV axis before P3 (yellow, BrdU-/EdU- cells) or after P3 (blue, EdU+ cells) in a single retina section (See Supplementary Fig. 3A-B for more examples). (**C**) A peak calling algorithm called local peaks of the SWA of the proportion of each subtype born before P3 (yellow curve in **(B)**, BrdU-/EdU- cells). Peaks are plotted based on their location along the DV axis arranged by subtype. (**D**) Loess-smoothed SWA of the local proportion of Type 6 bipolars in S-phase during the BrdU injection on P3. Curves show the local proportion of Type 6 cells in terminal (green = BrdU+/EdU- Type 6 cells), or non-terminal S-phase on P3 (red = BrdU+/EdU+ Type 6 cells) (See Supplementary Fig. 3G, and 3K-L for more examples). (**E**) A peak calling algorithm called local peaks of the SWA of the proportion of each subtype born on P3-P4. Peaks are plotted based on their location along the DV axis, arranged by subtype.

In addition to the consistent locations of peak early birthdates for different subtypes, the size of the alternating regions of enrichment and the magnitude of local enrichment were similar across subtypes and retinas. Consistently, peak widths covered roughly 10% of the retina arclength, with mean width of 10.57% ± 5.46% across all subtypes and retinas (Fig. 5D). Peak widths were consistent across retinas, and pair-wise t-tests revealed no significant differences between the distribution of peaks among all subtypes, across retinas (Supplementary Fig. 3D). Local peaks marked regions with approximately 10-30% enrichment in bipolar cells born, relative to adjacent trough regions (Supplementary Fig. 3E). Peak heights were similar in magnitude across retinas, but Retina 2 had significantly higher peak heights than Retina 1 (mean 29% enrichment and 20% enrichment, respectively). This difference could reflect slight differences in section location and angle across animal replicates or in precise developmental timing across animals, as different retina sections would intersect the underlying pattern at different spaces and times.

Taken together, we find that the retina is tiled with regions spanning 10% of the arclength that produce more bipolar cells during the first half of bipolar cell genesis (before P3) compared to directly adjacent regions. After P3, the regions of enriched bipolar cell birthdates swap: previous “troughs” of bipolar birthdates before P3 become “peaks” after P3, with more bipolar cell birthdates during the latter half of bipolar genesis. The location and size of these tiling regions, and the magnitude of the local enrichment of bipolar birthdates, are consistent across retinas and subtypes.

While these data suggest that birthdates of different subtypes are spatiotemporally correlated during the first and latter half of bipolar genesis, we asked how the subtype birthdates are patterned during peak bipolar genesis at finer time resolution. First, we considered the terminal S-phase population (BrdU+/EdU-), which includes cells born in the 16 hour window between P3 and P4. Birthdate modelling in Fig. 3 suggests that at this critical time, the OFF subtypes have shifted to their latter half of genesis, and the ON subtypes are still within their first half of genesis (Fig. 3D).

As a pooled population, bipolar cells born on P3-P4 were not patterned in peaks and troughs across the retina space (Supplementary Fig. 3F). However, individual subtypes display clear peak-trough patterns across all subtypes and retinas, with periodic hotspots of local subtype production (Fig. 5D green curve; (Supplementary Fig. 3G). The size of hotspots were consistent across retinas among all subtypes, or compared to the peak widths observed at coarser time resolution, before P3, covering an average of 9.83% ± 4.13% (mean ± std) of the retina arclength across subtypes and retinas (Supplementary Fig. 3H). Average peak height was 17% across subtypes and retinas, which is slightly but significantly lower than the 24% average peak height before P3 (*p* = 4.32 × 10^-8^). Peak heights across retinas were not significantly different (Supplementary Fig. 3I).

During this 16 hour window, peak locations were not consistent across all subtypes in the same retina (Fig. 5E, Supplementary Fig. 3G, J-K). These data demonstrate that bipolar cells born on P3-P4 are also patterned in alternating regions of enrichment along the retina space, in ‘hotspots’ of localized bipolar subtype production, but that hotspot locations are different for different subtypes.

To understand how the pattern changes in time, we examined birthdates and locations of cells that were in a non-terminal S-phase on P3. This population was in S-phase at the same time as the P3-P4 born population discussed above but went on to divide again rather than exit, producing bipolar subtypes during subsequent cell divisions. Across all subtypes and retinas, peaks and troughs of subtype birthdates were also observed among this P3 non-terminal S-phase population (examples in Fig. 5D, Supplementary Fig. 3L). Peak locations of the non-terminal S-phase population were significantly displaced from those on P3-P4, with a mean distance of 0.0618 ± 0.004 (mean ± s.e.m.). Since a distance of 0.05 arclengths would precisely mark trough regions of the P3-P4 birthdated bipolars, this indicates that the non-terminal S-phase cells on P3 produced bipolar subtypes later, in nearly opposite regions. In particular, bipolar birthdates cluster in alternating areas over time.

Our data show that bipolar subtype birthdates are patterned in hotspots, such that local production of subtypes alternates across space over time, peaking in different locations for different subtypes on P3-P4. Integrating birthdates of bipolar cells born in the first or latter half of bipolar genesis, however, yields consistent patterns among subtypes in the same retina.

### 2.5. Postnatal clones are biased against containing homotypic bipolar subtypes

One model that could explain the observed peaks of locally coordinated subtype birthdates is that subtypes arise from biased RPCs, which locally produce homotypic bipolar cells during the first postnatal week. To address this, we examined the bipolar subtype composition in postnatal retinal lineages and whether clones contain homotypic bipolar cells (Fig. 6A).

**Figure 6.**
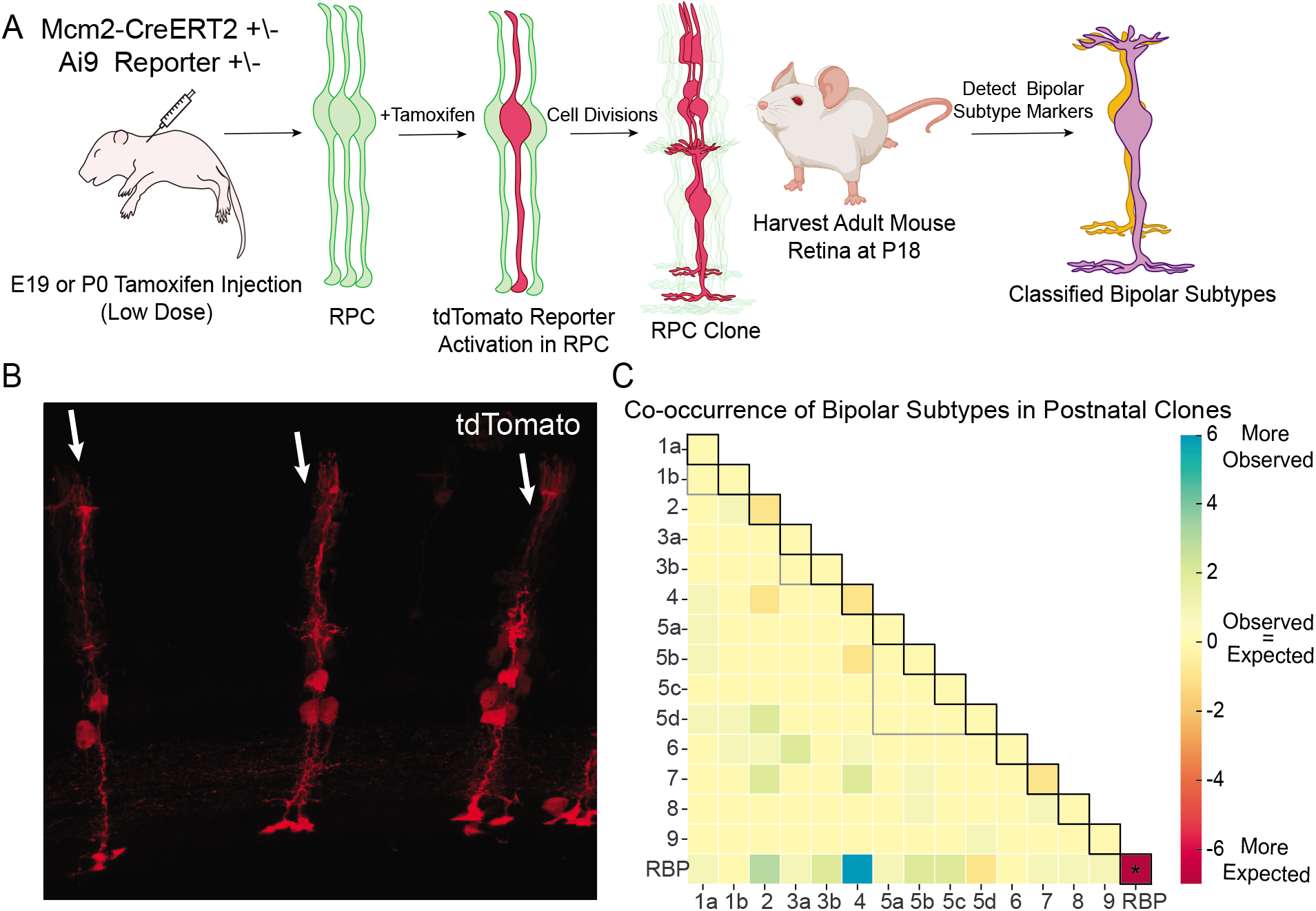
Late retinal clones are biased against containing homotypic bipolar subtypes. (**A**) Experimental schematic. Transgenic Mcm2-CreERT2 mice, with specific tamoxifen-inducible Cre expression in cycling cells, were crossed with Ai9 Cre reporter mice. Low doses of tamoxifen were injected at E19 or P0 to sparsely induce Cre recombination in isolated RPCs. Retinas developed *in vivo* until P18, at which point they were fixed, sectioned, and stained for bipolar subtype markers (described above) and the Cre reporter label. Isolated clones containing two bipolar cells were analyzed for their subtype composition. (**B**) Example image of postnatal retina clones labelled with tdTomato. Individual clones are marked with white arrows, and appear as radial columns of tightly associated cells. (**C**) Heatmap of pairwise clonal co-occurence of bipolar subtypes within postnatal clones containing pairs of bipolar cells (*n* = 82), plotted as enrichment relative to random expectation (see Methods). Monte Carlo analysis was performed to determine the expected frequency of pairwise co-occurrences of each subtype within clones. Colormap reflects the difference between observed number of clones and expected number of clones containing the subtype pair ([observed frequency - expected frequency]). Star indicates that the observed number of pair-containing clones was significantly different than the expected number (*p* < 0.05 after Bonferroni correction). Black outline shows pairs that contain the same subtype; grey outline shows pairings that include subtypes in the same class ({1a, 1b}, {3a, 3b},{5a, 5b, 5c, 5d}).

To label retinal clones, we took advantage of the progenitor-specific Cre line, Mcm2-CreERT2, where tamoxifen-inducible Cre is co-transcribed with a member of the DNA helicase complex [17]. *Mcm2* is specifically expressed in progenitor cells and is tightly regulated due to its role in DNA synthesis [18]. Mcm2-CreERT2 males were crossed to Ai9 reporter females, harboring a genomic, Cre-dependent tdTomato reporter. Delivery of low doses of tamoxifen to newborn mice or E19 pregnant females resulted in sparse labelling of clones, reflecting isolated reporter activation events in individual RPCs (Fig. 6B). Mice were harvested at P18, and retinas were fixed, sectioned, and stained for the 16 bipolar subtype markers, as described for the birthdating studies.

Since bipolar interneurons do not migrate from their clonal column [5,19,20], we inferred that radial columns with >50μm separation from other cellular columns were isolated clones deriving from a Cre recombination event in an isolated RPC. Radial columns of tightly packed cells containing rods, bipolars, amacrines, and Müller glia were labelled, consistent with previous reports of postnatal clone composition and morphology [20] (Fig. 6B). Furthermore, restricted reporter expression within postnatally-born cell populations and lack of reporter expression in earlier born cell types confirmed that the reporter activation was specific to mitotic cells.

With rare exception, clones with a bipolar cell contained one or two bipolar cells. Those with two bipolar cells were imaged for tdTomato expression (n = 82), along with SABER-FISH signals for all subtype markers. Bipolar subtypes were manually classified based on gene expression and morphologies to ensure correct identification and that clones were well-resolved. Surprisingly, we observed significant de-enrichment for clones containing homotypic bipolar cells and for clones containing same-class bipolar subtypes (Types 1a-b, 3a-b, 5a-d) (Fig. 6C). To statistically test whether the observed co-occurrence of subtypes in clones were different than the random expectation we built a Monte Carlo simulation to define the expected frequency of each subtype pair in clones (see Methods). The simulation predicted that 8 RBP/RBP pairs, based on the total number of RBPs analyzed. However, we observed only one such clone, demonstrating significantly fewer RBP pairs than expected (*p* = 0.0018). There were significantly fewer clones with the same subtype (*p* = 6.00 × 10 ^−5^) (Fig 6C, black outline), significantly fewer clones containing same-type cone bipolars (*p* = 0.01), and significantly fewer same-class bipolar subtypes than the random expectation (Types 1a-b, 3a-b, 5a-d; p = 1.00 × 10^-5^) (Fig 6C, grey outline). Interestingly, we observed RBPs in clones with every cone bipolar subtype, and although Type 4 and RBP pairs were observed more frequently than expected, the enrichment was not significant after correction for multiple hypothesis testing. These data suggest that the last few divisions of RPCs, which generate postnatal clones, are biased against producing homotypic bipolar cells. The observation that RBPs are present in clones with all cone bipolar subtypes and de-enriched for co-occurrence offers an explanation for the large number produced over similar developmental time (Fig. 3D): there are more RPCs that produce RBPs than other subtypes, rather than specialized RPCs that produce more RBPs.

### 2.6. A hierarchical model of neuronal subtype genesis

The current model of retinal development states that multipotent RPCs undergo changes in competence throughout time, gaining the ability to produce specific cell fates and losing the ability to produce others [21]. However, the emergence of subtype diversity within this framework has not been addressed, due to lack of tools.

Based on the ordered birthdates of bipolar subtypes and their wave-like arrangement across space, we hypothesized that they might emerge from a hierarchical lineage structure. The adult mouse retina is a mosaic of radial clonal units, each comprising the diversity of retinal cells that arise from individual, multipotent RPCs [5,20,22–24]. Proliferation occurs throughout the retina [14], and clonal analyses suggest that some terminally-dividing RPCs produce lineages with biased cellular compositions [25–28] but that clones arising from founding RPCs contain a grossly representative set of retinal diversity [24,29]. We noticed that the size of the peak regions approximates the size of radial clonal columns derived from the earliest marking of RPC lineages [24,29]. With the following assumptions, we built a model of the proliferating retina to test whether hierarchical production of bipolar subtypes and known proliferation dynamics could alone produce the observed wave-like birthdate patterns (Fig 7A) (see Methods).

**Figure 7.**
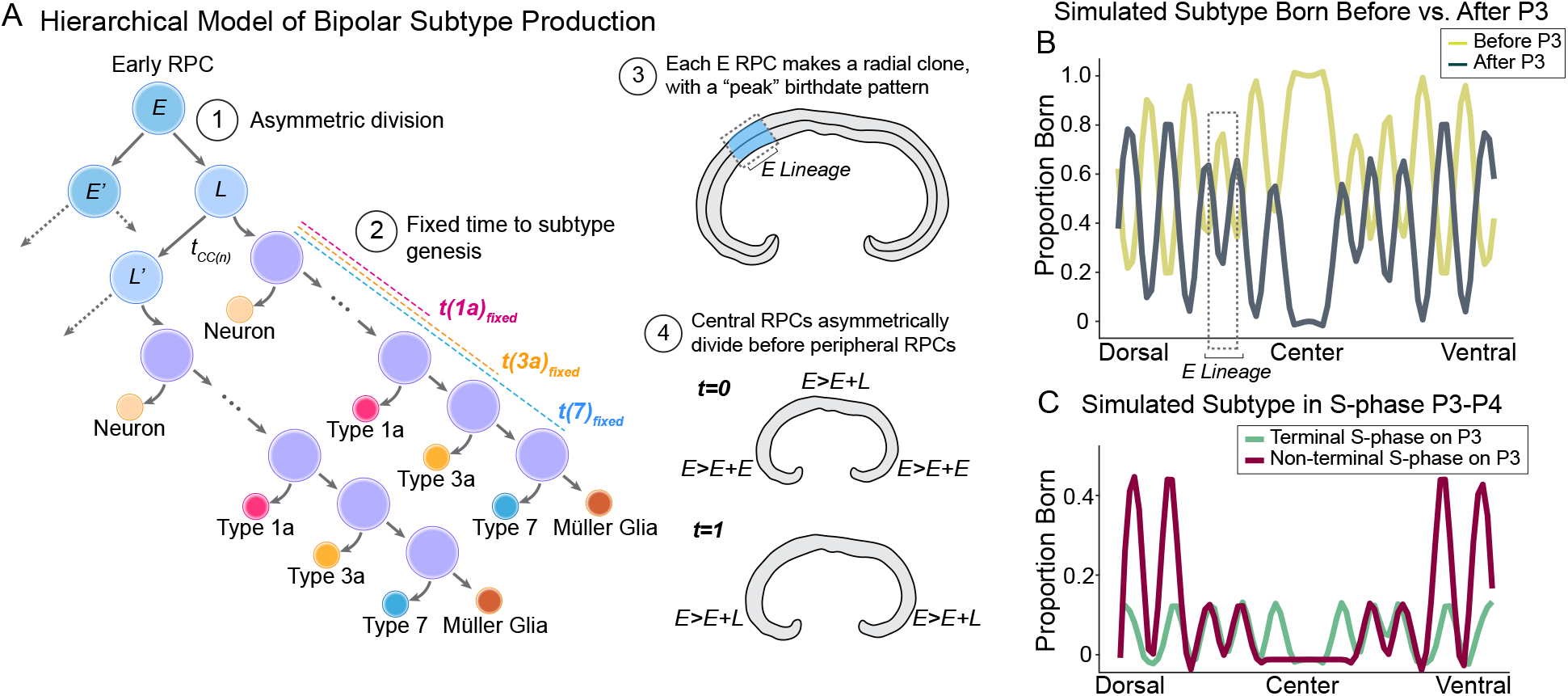
A hierarchical model of bipolar subtype genesis produces wave-like birthdate patterns. (**A**) A proposed and simulated hierarchical model of of bipolar subtype genesis. (1) RPCs in the early retina undergo symmetric divisions to expand the RPC pool (E>E+E). After a few rounds of division, RPCs divide asymmetrically (E>E’+L). L RPCs divide asymmetrically, and the non-L daughter produces neurons in a hierarchy, with a fixed time to subtype-specific birthdates before exiting mitosis to make Müller glia. (2) An example L lineage is shown, but the division pattern can vary and would include other retinal neurons, simply maintaining a fixed time to subtype genesis. *t*(*type*)_*fixed*_ indicates the fixed interval between L RPC genesis and subtype birthdate, which differs for different subtypes. (3) Cells deriving from each E lineage span large radial columns in the adult retina and display an internal peak birthdate distribution. The number of E cells sets the number of bipolar birthdate peaks/troughs across the retina, such that each peak corresponds to an E RPC and each trough marks the border of their clones with adjacent E lineages. (4) The shift from symmetric (E>E+E) to asymmetric (E>E’+L) divisions occurs in the central retina before the periphery. After the shift, the RPC progeny of L cells undergo fixed time to subtype genesis and mitotic exit. (**B**) Loess-smoothed SWA of the local proportion born before (yellow) versus after P3 (blue) of a simulated bipolar subtype, encoded by the model in **(A)**. (**C**) Loess-smoothed SWA of the local proportion in terminal (green) or non-terminal (red) S-phase on P3 for a simulated bipolar subtype, encoded by the model in **(A)**.

The early retina undergoes exponential growth, with founding RPCs symmetrically dividing between E9 and E11 before neurogenesis commences around E11.5 [30,31]. Therefore, our model begins with a slice of the early retinal epithelium, undergoing symmetric divisions (E>E+E) (Fig. 7A, (1)). Before the onset of neurogenesis, E RPCs in the central retina undergo an asymmetric division to self-renew and produce one intermediate L RPC (E>E+L) (Fig. 7A, (2)). At the onset of neurogenesis around E11.5, L RPCs begin to divide asymmetrically to generate retinal diversity and to generate more L RPCs. During this period, which we model to extend over 15 cell cycles in accordance with previous estimates [32], the bipolar subtypes are generated by L RPCs in a linear hierarchy, such that Subtype X is generated at time *t*, Subtype Y is generated at time t + 1, and so on, where t = 0 marks the time of initial asymmetric division (E>E+L) (Fig. 7A, (3)). Importantly, the precise pattern of divisions can vary, but the fixed time between initial asymmetric division (E>E+L) to subtype birthdate is crucial, as this event sets a series of delayed subclones that produce bipolar subtypes during sequential cell cycles. For example, if we consider the simplified lineage depicted in Fig. 7A, the two Type 1a bipolar cells differ in birthdate by a factor of one cell cycle, since one derives directly from the L RPC and the other derives from the L’ daughter RPC. Since the model assumes that the time from L RPC to Type 1a genesis is fixed (*t*(1*a*)_*fixed*_), then the birthdates differ by the length of the L>L’ asymmetric division, *t*_cc(*n*)_. Introducing a stochastic variable that permits sister cells to arrange as [S1 S2] or [S2 S1] with equal probability unbiases the direction of retinal local growth, consistent with known proliferation patterns and clonal expansion. Finally, the shift to asymmetric (E>E+L) division occurs in the central retina before the peripheral retina (Fig. 7A, (4)).

Several aspects of our observed subtype birthdate patterns are well-captured by this model. First, the peak-trough pattern emerges naturally, without additional constraints beyond an initial asymmetric division, and hierarchy of subtype production. Simulating measurements of bipolar subtype birthdates before versus after P3 (Fig. 7B) or on P3-P4 (Fig. 7C) show similar wave patterns to real data (compare to Fig. 5B,D). Simulating local birthdates before P3 for multiple subtypes results in correlated peak-trough patterns (Supplementary Fig. 4A), consistent with the real data (Supplementary Fig. 3B-C). At finer temporal resolution of birthdates on P3-P4 (Supplementary Fig. 4B), the patterns are not locally correlated among different simulated subtypes, consistent with real data (Supplementary Fig. 3G).

Although the encoded model captures key data features, it remains an oversimplification. For example, the peak heights of simulated subtypes exceed those in real data (compare Fig. 5B to Fig. 7B). Furthermore, from this simple model, it is unclear how small differences in average birthdates of bipolar subtypes can be observed, particularly those that are shifted by <1 cell cycle length (Fig. 3C). A possible extension of the model that could potentially resolve these discrepancies is the incorporation of multiple, asynchronous RPC populations that produce bipolar cells in parallel through the same hierarchical structure, but are shifted in time by a fraction of the cell cycle. Our encoded model has a single synchronized population of RPCs, but asynchronous RPC populations certainly do exist [33,34]. Further modelling could address these discrepancies, but the simple implementation demonstrates that a hierarchical lineage structure and passive expansion of the retina via local proliferation could alone generate the observed wave-like birthdate patterns.

## 3. Discussion

In this study, we constructed a comprehensive, high resolution map of the birthdate patterns of a diverse neuronal class. Using mesoscale multiplexed FISH to detect 16 RNA markers, we reconstructed full retina sections and identified all bipolar subtypes in a single experiment. Combining this with precisely timed injections of BrdU and EdU captured information about the time of mitotic exit, or cellular birthdate, of every cell. Automated analysis of the 3-D image data permitted investigation of spatial patterns of bipolar cell birthdates at scale, capturing nearly 10,000 single bipolar cells in each retina. This led to the discovery of previously unknown patterns of neuronal subtype genesis. Importantly, our methodology does not require automated machinery or a custom setup, and could be easily adapted to other experimental and biological systems.

### 3.1. Bipolar subtypes birthdates are ordered, reflecting their function

All 15 bipolar subtypes are born during the first postnatal week. By sampling nearly 30,000 bipolar cells and their birthdates, we achieve the statistical power to resolve their overlapping birthdates into subtype-specific birthdate patterns.

The data suggest that OFF subtypes are born before ON subtypes, and confirm that cone bipolar subtypes emerge before RBPs [12]. These results suggest that bipolar subtype birthdates correlate with visual function, and that subtype maturation follows the order of subtype births. Cone bipolar subtypes invaginate cone pedicles before RBPs invaginate rod spherules (reviewed in [35]), and the OFF sublaminae of the IPL matures before the ON sublamiae [36]. These observations suggest that the time of birth and maturation of bipolar subtypes are correlated, but further studies are required to track the process in individual cells.

The emergence of Types 4 and 1a as the first bipolar subtypes could also reflect a functional order. Type 1a laminates in the uppermost, scleral lamina of the IPL, while Type 4 laminates in the second sublaminae, demarcating the boundary between the OFF and ON layers [7,37]. It has been shown that the laminar layering of OFF versus ON bipolar cells happens normally in the absence of their ganglion cell afferents and subsets of amacrine cells within this layer [36,38] suggesting that the bipolar population might intrinsically carry out this process. It would be of interest to investigate whether earlier-born subtypes regulate lamination of later-born subtypes.

While many bipolar subtypes displayed significantly different birthdates, subtypes with indistinguishable birthdates are equally interesting. Types 2 and 3a, for example, had nearly identical average birthdates. Interestingly, loss of Irx6 in mice results in a hybrid bipolar cell where Type 3a bipolars acquire molecular and morphological features of both Type 3a and Type 2 [39]. To our knowledge, this is the only known mutation to produce a hybrid bipolar phenotype and the indistinguishable birthdates of Types 2 and 3a might reflect their origin from a common multipotent precursor cell.

### 3.2. Bipolar subtype genesis proceeds without the central retina

In both vertebrates and invertebrates, retina development is marked by a progressive front of differentiation that moves along a precise axis of the neuroepithelium. This pattern initially impelled a crystalline model, with early born cells inducing subsequent cell differentiation [40].

Birthdates of all bipolar subtypes follow a centrifugal pattern, consistent with previous observations [12]. This progression could permit early-born central bipolar cells to guide the production of later-born bipolar cells in the periphery. By removing the central retina on P0, we demonstrate that bipolar subtypes develop normally in the peripheral retina, without central cues. Our data support a model where chronology of bipolar subtype differentiation is intrinsically poised in the peripheral retina many days prior to mitotic exit and overt differentiation, contrary to an inductive model.

A similar experiment was performed on the firstborn neurons in the embryonic chick retina, retinal ganglion cells, finding that their timing of differentiation was intrinsic and respected the original developmental axis [41]. Together, these experiments suggest that the clock is intrinsically set for patterning prior to birthdates or overt differentiation and that retina epithelial history translates to orderly neuronal development. This, and complementary work, point to intrinsic control of orderly neurogenesis in space and time.

### 3.3. A proposed model of hierarchical bipolar subtype genesis

To date, few insights have been made into the logic of diverse subtype neurogenesis in the vertebrate central nervous system. Our data provide a window into this process, suggesting that production of bipolar subtypes alternates in space and time (Fig. 5).

We propose a simple model to explain the observed spatiotemporal patterns of bipolar subtype birthdates, consistent with previous observations of retinal lineages and proliferation. The model prescribes that 1) early RPCs divide asymmetrically, to produce hierarchical neurogenic lineages that are staggered in time, and 2) the number of total cell cycles, from the neuroepithelial stage RPC to Müller glia production, is relatively consistent throughout the retina, resulting in a retina that expands locally during the period of neurogenesis. From this model, it is easy to see that the number of peaks of local early birthdate enrichment equals the number of initial E RPCs (after the period of early symmetric expansion), and that the total cell number in the adult retina can be scaled by adding symmetric or amplifying divisions within the L lineages. The model does not require strict lineage patterns, consistent with previous characterization of clonal variability (examples in [19,20]), and fluidity in the composition of L lineages could generate the full diversity of retinal neurons. By invoking an internal hierarchy coupled with the simple fact that single RPCs give rise to two cells, the model demonstrates that wave-like birthdate patterns of neuronal subtypes would naturally arise.

Future refinement of the proposed model could incorporate heterogeneity in RPC populations [33,34], a probabilistic hierarchy for subtype production over time, and other aspects of known proliferation dynamics and cell death [42–44]. Evidence of biased terminal divisions exists [25,26,28], and it is possible that lineage biases also exist in the early retina as has been observed in *Xenopus* [45,46]. This factor could be incorporated into our model, by restricting populations of L lineages to produce specific types or subsets of types. In accordance with clonal data, we believe that the earliest E RPCs at the neuroepithelial stage give rise to clones with representative retinal diversity, as observed in [29]. After shifting to L lineages, biased clonal populations may emerge and clones could separate into disjoint radial columns as the retina grows [5,19,20,24]. Assuming that each peak is symmetric and corresponds to the clone of a founding RPC, we estimate that the retina derives from approximately 450 RPCs, which parallels the previous estimate of 500-1000 based on analyses of chimeric retinas [32]. Taken together, we believe that this model very simply captures the observed pattern of bipolar subtype birthdates in the retina with very few assumptions, and is consistent with previous clonal and proliferation data.

Importantly, the proposed model accounts for the most striking aspect of retinal development: the tiling of functional circuitry across the retina by local production of cellular diversity. While cellular migration and death contribute to the final distribution of some retinal cells [47], the production of diversity locally is a main organizing prinicpal of retinogenesis. The observation that late retinal clones do not contain homotypic bipolar subtypes is also consistent with this model and provides evidence for selective production of local diversity. Further studies are required to understand how homotypic restriction is imparted in postnatal clones, and increased n are required to assess preferential lineage patterns among subtypes.

It has been proposed that the RPC population is heterogeneous [21], but single-cell transcriptomics have not revealed obvious progenitor population distinctions that can explain the production of all retinal cell types [33,34]. The proposed model could explain the lack of obviously distinct RPCs, even if RPCs with deterministic lineages do exist. Our model suggests that RPCs in precise competence states would exist transiently along the retina space-time continuum as a tiny portion of the cellular population. This handful of synchronized cells would therefore be rare among the population sequenced and difficult to identify among transcriptional drop-outs and noise. Therefore, we believe that spatial analyses will be required to capture meaningful RPC differences that might reflect differential competence to produce neuronal subtypes, as we expect these populations might exist in spatial patterns at a given space and time might exist in predictable patterns at a given space and time.

How can this model be tested? An essential piece is to overlay clonal relationships with cellular birthdates to observe the order in which bipolar subtypes are generated by individual RPCs. For these future studies, capturing the full diversity of neuronal subtypes is essential for relative birthdate comparisons within single lineages. Here, we outline a method for identifying bipolar diversity in situ and established some ground-truth clonal relationships the subtypes (Fig. 6). Therefore, the bipolar system is an ideal application for emerging lineage tracing methods, which might be capable of capturing the fine structure of retinal lineages. Careful tracking of proliferation along the DV axis could also refine or refute the proposed model.

Other models could generate the observed birthdate pattern. Since our analyses were performed at P18 after bipolar cell death [42,43], we cannot infer its influence on the final pattern. We believe, however, that the pattern is unlikely to arise solely from cell death based on the amount and specificity of subtype death that would be required. It is also possible that the pattern is shaped or fine-tuned by real-time environmental feedback [11]. Examples of extrinsic signals influencing RPC fate decisions have been observed that fine-tune the proportions of RGCs and amacrine cells [48,49], but the impact of extrinsic cues on subtype frequencies has not been explored. The existence of such feedback is not inconsistent with this model and could act as a layer of regulation beyond intrinsic lineage structure for more robust development. Similar approaches applied in this manuscript could be overlayed with mutants that prevent apoptotic cell death [44] or disrupt extrinsic signaling to explore these contributions.

### 3.4. A window into the logic of neuronal subtype genesis

Our results focus on the development of bipolar subtype diversity, but we imagine that this model extends to other retinal cells. Since some classes of amacrine cells display ordered genesis [13,16], it is reasonable to consider that they too could emerge hierarchically within lineages. We believe that the proposed system would provide a robust developmental strategy to generate the balance of cellular diversity throughout the retina, relying solely on early asymmetric divisions, a simple intrinsic temporal axis [21], and passive proliferation to locally expand space as the retina develops.

## 4. Methods

### 4.1. CONTACT FOR REAGENT AND RESOURCE SHARING

Requests for resources and information should be directed to and will be fulfilled by the Lead Contact, Constance Cepko.

### 4.2. EXPERIMENTAL MODEL AND SUBJECT DETAILS

### 4.3. Tissue

All animal experiments were conducted in compliance with the protocol IS00001679, approved by the Institutional Care and Use Committees (IACUC) at Harvard University. Experiments were performed on tissue collected from wild-type male and female CD1 IGS mice (Charles River). Tissue were collected at postnatal day P18 for all birthdate experiments, and P17-P21 for all clone experiments.

### 4.4. RNA SABER-FISH probe design

A panel of bipolar subtype markers was selected based on subtype-specific gene expression profiles described in [4]. These markers were selected based on specificity and level of expression in subsets of bipolar subtypes.

SABER-FISH probe sets were designed to target each marker mRNA as previously described[6], using the Oligominer pipeline [50] (source code available at https://github.com/beliveau-lab/OligoMiner). Probe sets were ordered for each marker gene, with a unique SABER primer sequence for each gene permitting independent detection in the same tissue. Primer sequences 25-28, 30-34, 36-37, 39, and 41-44 were used, taken from [6]. Probes for each gene were taken from the precompiled mm10 whole-genome probe set, hosted at (https://oligopaints.hms.harvard.edu/genome-files).

### 4.5. Nucleoside analog injections for ‘window labelling’

Mice were injected subcutaneously on P3 at 17:00hr with 50ul of 2.5mM BrdU in PBS (ThermoFisher Scientific, B23151). Sixteen hours later, they were injected with 50ul of 1mM EdU in PBS (ThermoFisher Scientific, Click-iT^™^ EdU Cell Proliferation Kit for Imaging, C10340) and every 12 hours through P7. The EdU 12-hour injection times were chosen based on known cell cycle kinetics [14] to label all cells cycling after P4 (Supplementary Fig. 2A-B).

### 4.6. Tissue preparation

Mice were harvested at P18 and retinas were prepared for analysis as previously described [6]. Briefly, retinas were dissected in PBS and fixed for 30 minutes in 4% paraformaldehyde (v/v), 0.25% Triton-X (v/v). After fixation, retinas were washed 2 times in PBS for 5 minutes, transferred to 7% sucrose in PBS (w/v) for 10 minutes, and transferred to a 1:1 solution of [30% sucrose in PBS]:[OCT] for 1 hour. Retinas were then frozen and stored at −80°C.

### 4.7. SABER-FISH for RNA marker detection

Ibidi 8-well Poly-L-Lysine coated chamber slides (Ibidi, 80826) were coated with Poly-D-Lysine overnight (0.3 mg/ml in 2X Borate Buffer) and rinsed with water in the morning. For each of three retinas, three *35μm* retina cryosections were collected from the central retina and placed into the chamber slide. Sections were washed 3 × 5 minutes with PBS+0.1% Tween (PBSTw). For clone experiments shown in Fig. 6, the endogenous tdTomato reporter was bleached by incubating the tissue in 100% methanol for 1 hour. Retinas were subjected to a 20 minute gradient to ease the methanol stansition, where they were incubated in 20%, 40%, 60% and 80% methonol (in PBSTw) solutions. Sections were then incubated with 100% methanol for 1 hour, followed by the reverse 20 minute gradient to return to PBSTw. For all experiments, retinas were moved to 43°C in Wash Hyb Solution (40% formamide, 0.1% Tween in 2XSSC) for 30 minutes.

SABER-FISH protocol was conducted on retina sections as published in [6]. For staining cell membranes, WGA conjugated to 405s (Biotium, 29027) was diluted to a concentration of 10 μg / mL in PBSTw and samples were incubated for 15 minutes at 37°C during the SABER fluorescent detection step. Slides were washed 2 X 5 minutes in PBSTw following WGA application. All imaging was done for SABER-FISH marker gene detection before other histological stains, and each imaging session contained the WGA counterstain in the 405 channel for alignment across sessions.

### 4.8. Detection of the tdTomato reporter for clonal analysis

Since the tdTomato signal was bleached for SABERFISH RNA detection, it was detected by immunohistochemistry after all RNA detection was complete and all fluorescent imager oligonucleotides were removed. A polyclonal rabbit anti-mCherry antobody was used (Abcam, ab167453), followed by a secondary Donkey anti Rabbit fluorescent antobody (Jackson Immunoresearch, AffiniPure Donkey Anti-Rabbit IgG (H+L)).

### 4.9. Microscopy

Samples were imaged with a Yokogawa CSU-W1 single disk (50 μm pinhole size) spinning disk confocal unit attached to a fully motorized Nikon Ti2 inverted microscope equipped with a Nikon linear-encoded motorized stage with a Mad City Labs 500 μm range Nano-Drive Z piezo insert, an Andor Zyla 4.2 plus (6.5 μm photodiode size) sCMOS camera using a Nikon Apo S LWD 40X/1.1 DICN2 water immersion objective lens with Zeiss Immersol W 2010. The final digital resolution of the image was 0.16 μm/pixel. Fluorescence from (405, 488, 550, and 640) was collected by illuminating the sample with directly modulated solid-state lasers 405 nm diode 100mW (at the fiber tip) laser line, 488 nm diode 100 mW laser line, 561 nm DPSS 100mW laser line and 640 nm diode 70mW laser line in a Toptica iChrome MLE laser combiner, respectively. A hard-coated Semrock Di01-T405/488/568/647 multi-bandpass dichroic mirror was used for all channels. Signal from each channel was acquired sequentially with hard-coated Chroma ET455/50, Chroma ET525/36 nm, Chroma ET605/52 nm emission filters and Chroma ET705/72 nm in a filter wheel placed within the scan unit, for blue, green, red and far-red channels, respectively. All images were captured using the 16bit dual gain high dynamic range camera mode and no binning. Nikon Elements AR 5.02 acquisition software was used to acquire the data. Z-stacks were acquired using Piezo Z-device, with the shutter closed during axial movement. Images were acquired by collecting the entire Z-stack in each color. Data were saved as ND2 files.

### 4.10. Image processing

For each region imaged, a total of 7 sequential multichannel images were acquired, each including a subset of bipolar subtype markers and a WGA counterstain. The consistent WGA counterstain was used to register images from multiple sessions for pixel-level alignmment to the first session, by the *antsRegistrationSyNQuick* function from the *Advanced Normalization Tools* package (http://stnava.github.io/ANTs/) that employs a series of rigid, affine, and diffeomorphic registrations [51]. All alignments were manually checked, and regions where any of the 7 images failed to align were excluded from analysis. Raw data and aligned images are available upon request.

### 4.11. Segmentation of cells, SABER-FISH signals, and nucleotide analog stains

Segmentation of retina cells in 3-D was performed using a published algorithm, *ACME* [52]. The WGA counterstain serves as a cellular outline, and the WGA stain from the first image session was used to generate a watershed segmentation of the cells in three dimensions.

SABER-FISH signals were segmented in 3-D using previously published MATLAB code [6]. Briefly, the SABER-FISH images were filtered with a top hat filter to remove background and a 3-D Laplacian of Gaussian filter was applied to detect individual SABERFISH puncta. The threshold for puncta detection was manually checked for each channel of all images, to ensure proper signal segmentation.

EdU and BrdU labels were segmented to generate a binary mask for each label. Each image was manually adjusted for high contrast in ImageJ using *Image* → *Adjust* → *Brightness/Contrast* by adjusting the minimum and maximum intensity values. The adjusted images were used as input to MATLAB’s *imbinarize* function, which performs adaptive, local thresholding (see Supplementary Fig. 2C for segmentation results). The resulting binary mask of each nucleotide label was overlayed with cell segmentation, and cells were deemed EdU^+^ if 10% of their pixels overlapped with the EdU mask, and BrdU^+^ if 20% of their pixels overlapped with the BrdU mask.

The OPL was manually traced in ImageJ for each image region based on the WGA stain, to enable calculations of cellular location along the dorso-ventral axis and distances of cell bodies from the OPL. Final retina reconstructions were made by connecting the OPL from adjacent images.

### 4.12. Identification of bipolar subtypes in situ

SABER-FISH marker data was used to determine bipolar subtype identities of segmented cells. Segmented objects were filtered for volume, intensity in the WGA channel (since non-cellular objects might have high WGA signal), distance from the OPL, and positivity for the pan-bipolar marker, Vsx2 (>4 puncta captured in the cell). A gene expression vector was computed for each cell, with the number of SABER-FISH puncta detected for each marker gene and the integrated intensity value of each marker channel within that cell, both normalized across all cells. We found that by including both SABER-FISH puncta counta and integrated intensity rather than either one alone, the bipolar subtypes were better resolved on a dimensional reduction and when used as training data for a classifier. Each cell’s boundary was then eroded by 3 pixels, three times, and the internal puncta and intensity counts were recomputed. The eroded measurements were added to the gene expression vector for each cell, to help correct for any puncta located at cell-cell boundaries that might be misassigned in the uneroded measurement.

The tSNE algorithm was used to perform dimensional reduction on the resulting gene expression matrix for all cells and marker genes in Retina 1. Bipolar subtype clusters were identified manually in this first retina, based on the expression of subtype-specific marker genes among the clusters (Supplementary Fig. 1B-C). K-means clustering was used to parse cells into rough clusters, and individual cells within k-means clusters were examined in the raw data to validate subtype identities (Supplementary Fig. 1B). Types 8 and 1b were manually split from a single k-mean cluster, but were well-resolved at finer resolution based on expression of Grm6 in Type 8. Many example cells from each cluster were spot-checked by looking at the SABER-FISH data.

Identified cells from Retina 1 were used to train a random forest classifier to identify bipolar subtypes based on the marker gene expression matrix of SABER-FISH data, described above. Specifically, the SABER-FISH puncta counts and integrated intensity values (with erosions) for 14,534 cells were used to train a random forest classifier with 1,200 trees, using Python’s *sklearn* package. When tested on the remaining 1,000 identified cells, the classifier performed with unweighted mean f1-score per label of 0.91, and weighted mean f1-score of 0.93.

This classifier was directly applied to the SABERFISH marker gene expression matrices for all segmented cells in the other retinas, to determine the bipolar subtype identities of each imaged cell.

### 4.13. Statistical modelling of subtype birthdates over developmental time

To generate birthdate probability curves in Fig. 3D and estimates of average subtype birthdates in Fig. 3C, cells were binned into three categories based on their nucleoside analog labelling, as shown in (Fig. 3A), corresponding to cellular birthdates of P0-P3 (BrdU-/EdU-), P3-P4 (BrdU+/EdU-), and P4-P7 (BrdU-/EdU+). To convert these data into an estimated distribution of birthdates over developmental time, the bin counts were used as measurements of bipolar subtype genesis over time. To estimate the underlying distribution, it was assumed that the probability density function (PDF) of genesis for each subtype follows a normal distribution between P0 and P7. A grid-search algorithm was executed to find the optimal solutions 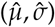 of the parameters (*μ, σ*) that minimized the difference between the observed bin counts and the estimated cumulative distribution function. Under these assumptions, the estimated distribution (PDF) for each subtype reflects the probability of generating that subtype at each day during development. In Figure 3C, the estimated average birthdate for each subtype was taken to be 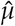 and the estimated standard error of the mean was computed as 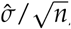, where *n* is the number of each subtype analyzed.

The proportion of each subtype born before P4 (BrdU-/EdU- and BrdU+ EdU-) versus after P4 (BrdU+/EdU+ and BrdU-/EdU+) was compared for each subtype pair using a binomial *z*-test. Bonferroni correction was performed to account for multiple hypothesis testing, and *p* values <0.05 after Bonferroni correction were considered significant (Fig 3E).

### 4.14. Central versus peripheral bipolar cell birthdates

The retina was divided into 13 regions and the proprotion of each bipolar subtype born before P3 (BrdU- /EdU-) was computed within each region. Regions 5-8 were used to calculate the average central proportion born, and Regions 1, 2, 12, and 13 were used to compute the peripheral proportion born for each subtype.

### 4.15. Retina explants

Retinas were explanted from P0 mice, right after birth. Retinas were dissected in 1X HBSS and quickly transferred to explant media, as defined in [**?**]. Briefly, the media is 50% Minimum Essential Medium (Millipore Sigma, 51412C)(v/v), 25% HBSS (Thermo Fisher Scientific, 14-025-092) (v/v), 25% heat-inactivated horse serum (Thermo Fisher Scientific, 26050-088), 200 μm L-glutamine (Sigma-Aldrich G3126), and 5.75mg/mL glucose, 25mM Hepes, 100 U/mL penicillin and 100ug/mL streptomycin (Invitrogen/Gibco 15140-122).

Centers were removed upon dissection using a 1mm biposy punch (World Precision Instruments, 504646). Retinas were then transferred to a 6-well plate, one retina per well. Each well contained 4mL of hardened 4% low-melt agarose (w/v), dissolved in the explant media. Retinas were pinned into the agarose, RGC layer facing up, using two insect pins. Retinas were covered with 1.5mL of explant media, and incubated at 34°C, 5% CO_2_ for 14 days. Each day, 1mL of media was replaced with new media.

### 4.16. Spatial analysis of bipolar birthdates

A continuous reconstruction was made of each sectioned retina from three sequential 35μm sections. Approximately 13 image regions at 40X magnification were acquired to cover each section. The location of each cell along the retina arclength was computed based on a manual trace of the OPL within each section, and this arclength was used for spatial analyses. In some cases, image regions were eliminated because at least one image session failed to align to the others. Even with these cases, however, the overlap of sequential sections resulted in full reconstructions for all retinas along the DV axis.

The average bipolar cell density was computed across the DV axis as a sliding window average (SWA), with window size of 10% of the retina arclength, and the average was computed in 100,000 iterations along the arclength. Similarly, a sliding window average of the proportion of each bipolar subtype born before P3, on P3-P4, and after P3 were computed with a window size of 5%, at 100,000 iterations along the arclength.

For visualization in (Fig. 5B, D, Supplementary Fig. 3A-B, G, K-L) Loess smoothing of SWAs was performed sampling 10% of the data points. Peaks were called in (Fig. 5C, E) by MATLAB’s *findpeaks* function based on the Loess-smoothed SWAs for each subtype in the single retina section highlighted in Fig. 5, with a ‘*MinPeakProminence*’ set to 0.01.

Pair-wise Pearson correlations were performed to compare the SWAs for subtypes in the same retina born before P3 (Supplementary Fig. 3C) or born on P3-P4 (Supplementary Fig. 3J). The numerical derivative of SWAs were also computed, to capture the local rate of change of the proportion of each subtype born across space, during each labelled time (before P3 or on P3-P4). Pearson correlations were computed pairwise for the derivatives, to assay whether the local areas of increased and decreased subtype birthdates were consistent among subtypes in the same retina. The pair-wise comparisons were done internally for each retina, and heatmaps display the average correlation across three retina replicates. (Supplementary Fig. 3C and 3J, right)

### 4.17. Toy model of hierarchical bipolar subtype genesis

The developing retina at a given time point is modelled as an one-dimensional array *X_t_*, where the position of an element corresponds to the cellular (and later clonal column) position along the dorsoventral arclength of a retina, and the element values representing the current cellular state. Elements of array *X_t_* can take a state among {*E_i_, L_i_, N_i_, M*}, where the subscript *i* maintains the number of cell divisions relative to the initial state transition. For simplicity, starting from an array of *E* state progenitors, the array is updated and expanded based on mapping rules in discrete time steps in synchrony, until no further updates are necessary. Mapping rule for each element in *X_t_* to *X*_*t*+1_ is set as follows:

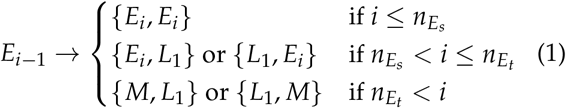

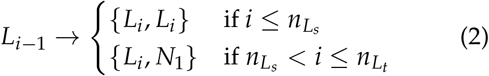

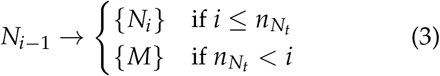

where *n_E_s__, n_L_s__* denote the number of cell cycles before switching to asymmetric division and *n_E_t__, n_N_t__* denote the last cell division before transitioning into terminal *M* (Müller glia) state for *E*, and *N* cell state respectively.

The *N* state cell is modelled to generate a specific bipolar subtype neuron for each time step (cell cycle), in a defined temporal order at the same cellular position such that eventually a clonal column is generated. As a result, all neurons have defined DV position (array index) as well as birth time (*t* when born) for analysis. Essentially, Eq. (3) models an asymmetric division tree generating a series of subtypes of neurons for each divisions. As long as the birth order of subtypes is maintained, any hierarchical tree representation can replace Eq. (3) to model the spatial wave pattern. The central-to-peripheral difference is incorporated in the model by setting the initial *E* cell cycle number *i* of the center to an advanced number than the periphery. The parameter values for Fig. 7 is as follows: *n_E_s__* = 2, *n_L_s__* = 2, *n_E_t__* = 16, *n_N_t__* = 4.

### 4.18. Clonal labelling

To label retinal clones, Mcm2-CreERT2 male mice were crossed with homozygous Ai9 Cre-reporter female mice. Plugs were checked to determine E0. For labelling of E18 and E19 clones, pregnant E18 or E19 female mice were intraperitoneally injected with 50ul of 2mg/ml of tamoxifen (Sigma-Aldrich, T5648) dissolved in corn oil. Late on E19, a cesarean section was performed to remove the pups, to prevent mothers from eating them upon birth. For clonal labelling on P0, pups were injected with 30-50ul of 0.05-0.1mg/mL of tamoxifen dissolved in corn oil. Injection volumes are difficult to control, especially into P0 pups, and result in variable clone densities. Retinas with sparse clone labelling were chosen for analysis.

### 4.19. Monte Carlo simulation for defining a random expectation of clonal compositions

To simulate the random expectation of bipolar subtypes within clones, we built a Monte Carlo simulation. We pooled all individual bipolar cells that were analyzed in clones containing two bipolar cells. A few clones containing 3 bipolar cells were analyzed, but these were omitted from the formal analysis for simplicity. The clones observed with three bipolar cells were: [Type 2, 5d, 7], [Type 8, Type 7 Type 5b], [RBP, Type 4, Type 5d], [RBP, Type 2, RBP].

We pooled all bipolar cells observed in the clones (which included all subtypes) and randomly drew 82 pairs from the set without replacement 100,000 times. For each pair of subtypes, the mode of the frequency of that pair in the 82 trials was taken as the expected number of clonal co-occurrences. The *p*-value was derived from the empirical cumulative distribution function of the Monte Carlo simulation, and *p*<0.05 after Bonferroni correction for multiple hypothesis testing was considered significant.

## Author Contributions

Conceptualization, E.W., S.L., and C.C.; methodology, E.W., S.L.; software, E.W.,C.L.; validation, E.W., S.L.; formal analysis, E.W., X.L.; investigation, E.W., S.L.; resources, E.W., S.L., K.K.; data curation, E.W.; writing—original draft preparation, E.W.; writing—review and editing, E.W., C.L., K.K., C.C.; visualization, E.W., C.L.; supervision, C.C.; funding acquisition, C.C.

## Data Availability Statement

Raw and processed data as well as scripts to generate plots or perform analyses are available upon request. Code will be published at (https://github.com/ewest11).

## Acknowledgments

The authors would like to thank Paula Montero-Llopis, Ryan Stephansky, and Timothy Ross-Elliot of the Harvard MicRoN core for helpful discussions, training, and collaboration. We thank Steven Pruitt for generously sharing the Mcm2CreERT2 line with us. We thank Grace Wallick, a technician in the Cepko Lab, for her technical help with mouse genotyping. We thank Emma’s miniature dachshund, Beans, for his expertise on retinal development and help editing the manuscript. This PDF preprint was generated from a LaTeX template provided by *MDPI* at (https://www.mdpi.com/authors/latex).

