## Supplementary Materials for "Spatiotemporal patterns of neuronal subtype genesis suggest hierarchical development of retinal diversity"

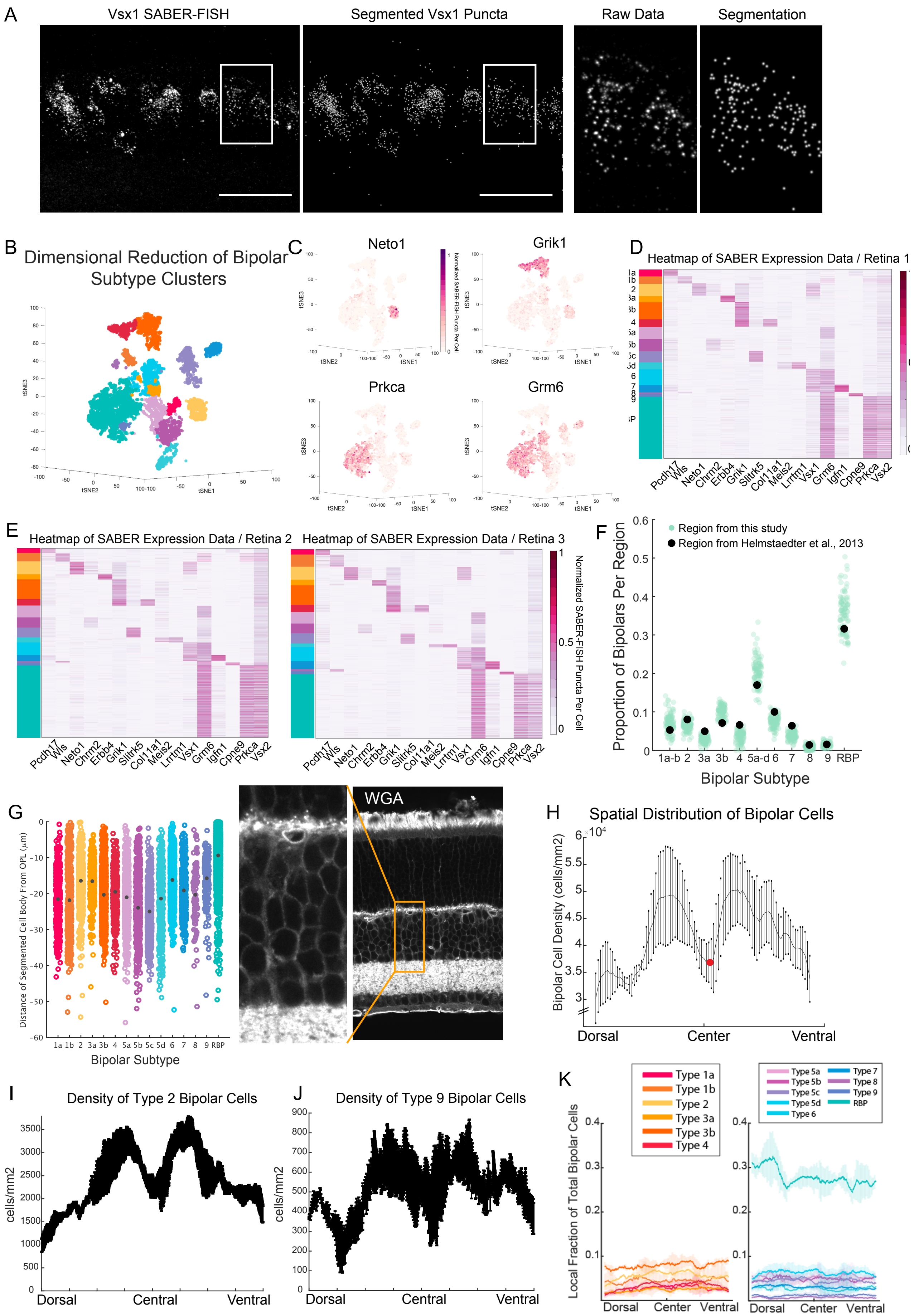
A
Vsx1 SABER-FISH
Segmented Vsx1 Puncta
Raw Data
Segmentation
B
Dimensional Reduction of Bipolar Subtype Clusters
C
Neto1
Grik1
Prkca
Grm6
D
Heatmap of SABER Expression Data / Retina 1
E
Heatmap of SABER Expression Data / Retina 2
Heatmap of SABER Expression Data / Retina 3
Normalized SABER-FISH Puncta Per Cell
Pcdh17 Wls Neto1 Chrm2 Erbb4 Grik1 Slitrk5 Col11a1 Meis2 Lrrtm1 Vsx1 Grm6 Igfn1 Cpne9 Prkca Vsx2
F
Region from this study
Region from Helmstaedter et al., 2013
Proportion of Bipolars Per Region
Bipolar Subtype
1a-b 2 3a 3b 4 5a-d 6 7 8 9 RBP
G
Distance of Segmented Cell Body From OPL (μm)
1a 1b 2 3a 3b 4 5a 5b 5c 5d 6 7 8 9 RBP
Bipolar Subtype
WGA
H
Spatial Distribution of Bipolar Cells
Bipolar Cell Density (cells/mm2)
Dorsal
Center
Ventral
I
Density of Type 2 Bipolar Cells
cells/mm2
Dorsal
Central
Ventral
J
Density of Type 9 Bipolar Cells
cells/mm2
Dorsal
Central
Ventral
K
Type 1a
Type 1b
Type 2
Type 3a
Type 3b
Type 4
Type 5a
Type 5b
Type 5c
Type 5d
Type 6
Type 7
Type 8
Type 9
RBP
Local Fraction of Total Bipolar Cells
Dorsal
Center
Ventral

**Figure S1. Identification of all bipolar subtypes *in situ*.** (A) **(A)** SABER-FISH puncta segmentation for bipolar subtype markers was done using a 3-D Laplacian of Gaussian method. Maximum intensity projections of raw SABER-FISH image of Vsx1 detection are shown (left) next to segmented puncta (right). Far right shows magnification of outlined region. Scale bars are 50µm. **(B)** Puncta counted in **(A)** were assigned to individual imaged cells based on 3-D cell segmentation (see Methods). All cells Vsx2+ cells in a single retina were pooled, and the tSNE algorithm was used to reduce the dimension of SABER-FISH single-cell data to two dimensions. Clusters were identified (left) based on subtype-specific gene expression patterns (right). Neto1 and Grik1 mark Type 2 bipolar cells, while Prkca and Grm6 mark RBPs, for example. Colormap corresponds to the number of SABER-FISH puncta detected per cell, normalized across all cells. **(D)** Heatmaps of SABER-FISH puncta per cell quantified for bipolar subtype markers for Retina 1. Colormap corresponds to the number of puncta per cell for each marker, normalized across all bipolar cells. Rows of the heatmap correspond to single cells, sorted by their assigned subtype identity. **(E)** A random forest classifier was trained on 14,534 cells from Retina 1 to identify subtypes based on SABER marker gene expression data in other retinas. Heatmaps of SABER-FISH marker gene expression across cells are shown for Retinas 2 (left) and 3 (right), sorted by subtype identities that were determined by the classifier. Colormap on the left corresponds to the same subtypes labelled in **(C)**. **(F)** Subtype frequencies were calculated as a proportion of all bipolar cells in each image region across 3 retinas (green dots). For comparison, the proportion of each subtype calculated in (Helmsteadter et al., 2013) based on EM reconstructions is shown (black dots). **(G)** Cell body locations of classified bipolar subtypes, plotted as distance relative to the OPL. Data reflect bipolar cells from Retina 1, including all cells in **(H)** The sliding window average (SWA) of measured bipolar cell density along the retina center-line (cells/mm$^2$), plotted across the DV axis. The line is the average regional density across 9 DV sections from 3 retinas. Error bars show standard deviation across retinas and red dot marks the location of the optic nerve head. **(I)** Sliding window average (SWA) of the density of Type 2 bipolar cells across the DV axis along the retina center-line. Error bars reflect standard deviation across three retina replicates. **(J)** Sliding window average (SWA) of the density of Type 9 bipolar cells across the DV axis along the retina center-line. Error bars reflect standard deviation across three retina replicates. **(K)** SWA of the local proportion of each subtype, as a fraction of the total bipolar population. OFF subtypes are shown on the left, and ON subtypes on the right. Shaded region represents standard deviation across three retina replicates.

A

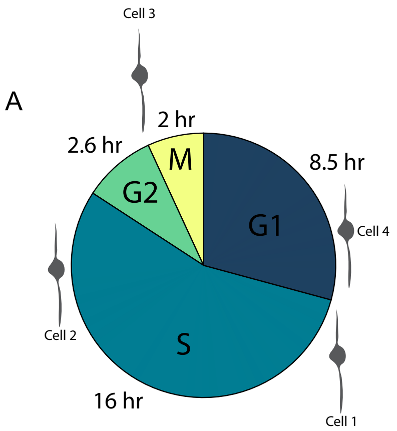


B

| | Status on P3 (time of BrdU injection) | Status 16 hours later (time of first EdU injection) | If it re-enters the cell cycle | If it exits after M phase |
|---|---|---|---|---|
| Cell 1 | Early S phase (BrdU+) | G2 (EdU-) | 12hrs later = S phase (EdU+) | 12hrs later = G0 (EdU-) |
| Cell 2 | Late S phase (BrdU+) | Depends on cell cycle reentry/exit | 16hrs later = S phase (EdU+) | 16hrs later = G0 (EdU-) |
| Cell 3 | G2/M/Early G1 (BrdU-) | Depends on cell cycle reentry/exit | 16hrs later = S phase (EdU+) | 16hrs later = G0 (EdU-) |
| Cell 4 | Late G1 (BrdU-) | Depends on cell cycle reentry/exit | 16hrs later = S phase (EdU+) | 16hrs later = G0 (EdU-) |

C

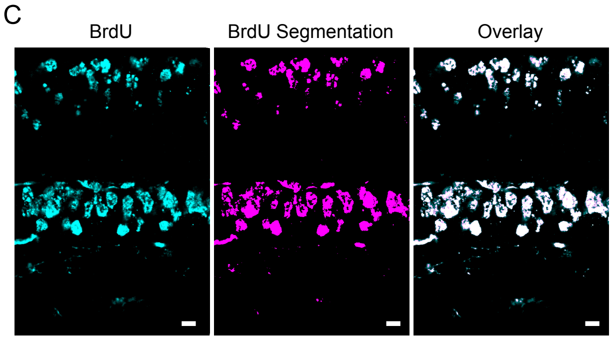


D

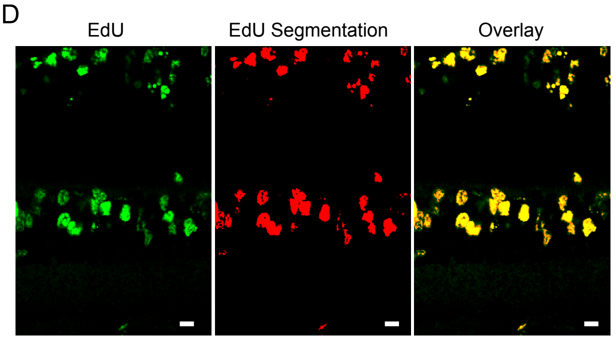


E

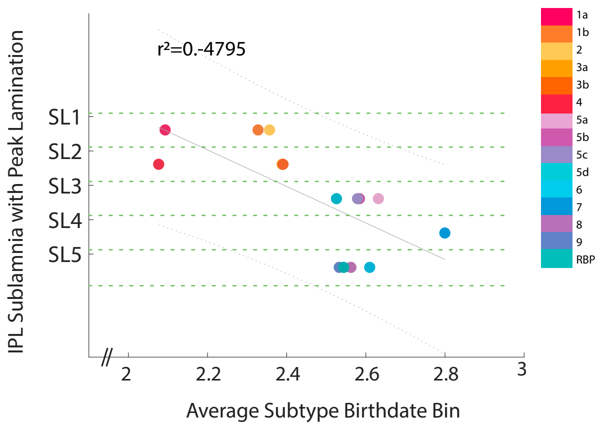


F

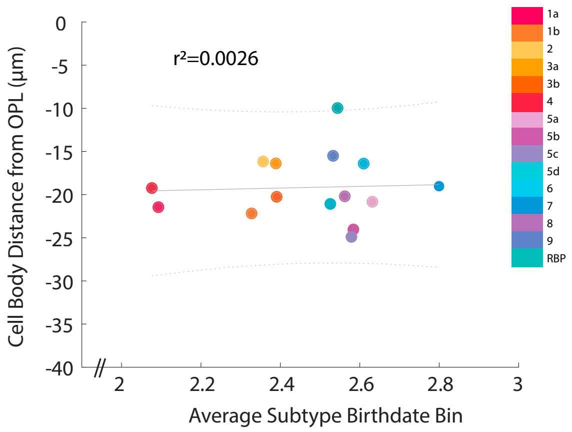

**Figure S2. Window labelling with BrdU and EdU allows precise cellular birthdating (A)** Schematic of the cell cycle, measured in postnatal day 1 mice in [1]. **(B)** Example cells in 4 different cell cycle phases at the time of BrdU injection on P3. Cells in S phase on P3 at 17:00hr are labelled with BrdU. If they reenter the cell cycle (Cell 1 or Cell 2), they will be labelled with EdU by the injection 12 hours later, at 9:00hr on P4. If an S phase cell does not reenter the cell cycle, then it will be BrdU+/EdU- and thus be "birthdated" within the P3 17:00hr and P4 9:00hr window. Cells in G2/M/G1 will not be labelled by the BrdU injection, but will be labelled by subsequent EdU injections if they re-enter the cell cycle after M phase (Cell 3 and 4). **(C)** BrdU signal detected by immunohistochemistry (left) was segmented to create a binary mask (center). Overlay of raw signal from a single z-plane with the segmented mask is shown (right). Cells were deemed $BrdU^{+}$ if more than 20% of their volume was positive for the BrdU segmented mask. Scale bar is 10$\mu$m. **(D)** EdU signal detected by click chemistry (left) was segmented to create a binary mask (center). Overlay of raw signal from a single z-plane with the segmented mask is shown (right). Cells were deemed $EdU^{+}$ if more than 10% of their volume was positive for the EdU segmented mask. Scale bars are 10$\mu$m. **(E)** Bipolar subtypes laminate in specific layers of the innerplexiform layer (IPL), as demonstrated by electron microscopy in [2]. Plot displays the IPL sumblinae of peak lamination versus the average birthdate bin for each subtype. Birthdate bins correspond to: 1=born before P3, 2=born on P3-P4, and 3=born after P3. Dotted lines mark IPL sublaminae (SL) 1-5, marked by 25% increments of IPL depth. Regression line is shown, with $r^2 = -0.4795$. **(F)** The distance of each segmented cell body from the OPL was calculated and plotted against the average subtype birthdates (unit explained in **(E)**). Regression line is plotted, with $r^2 = 0.0026$.

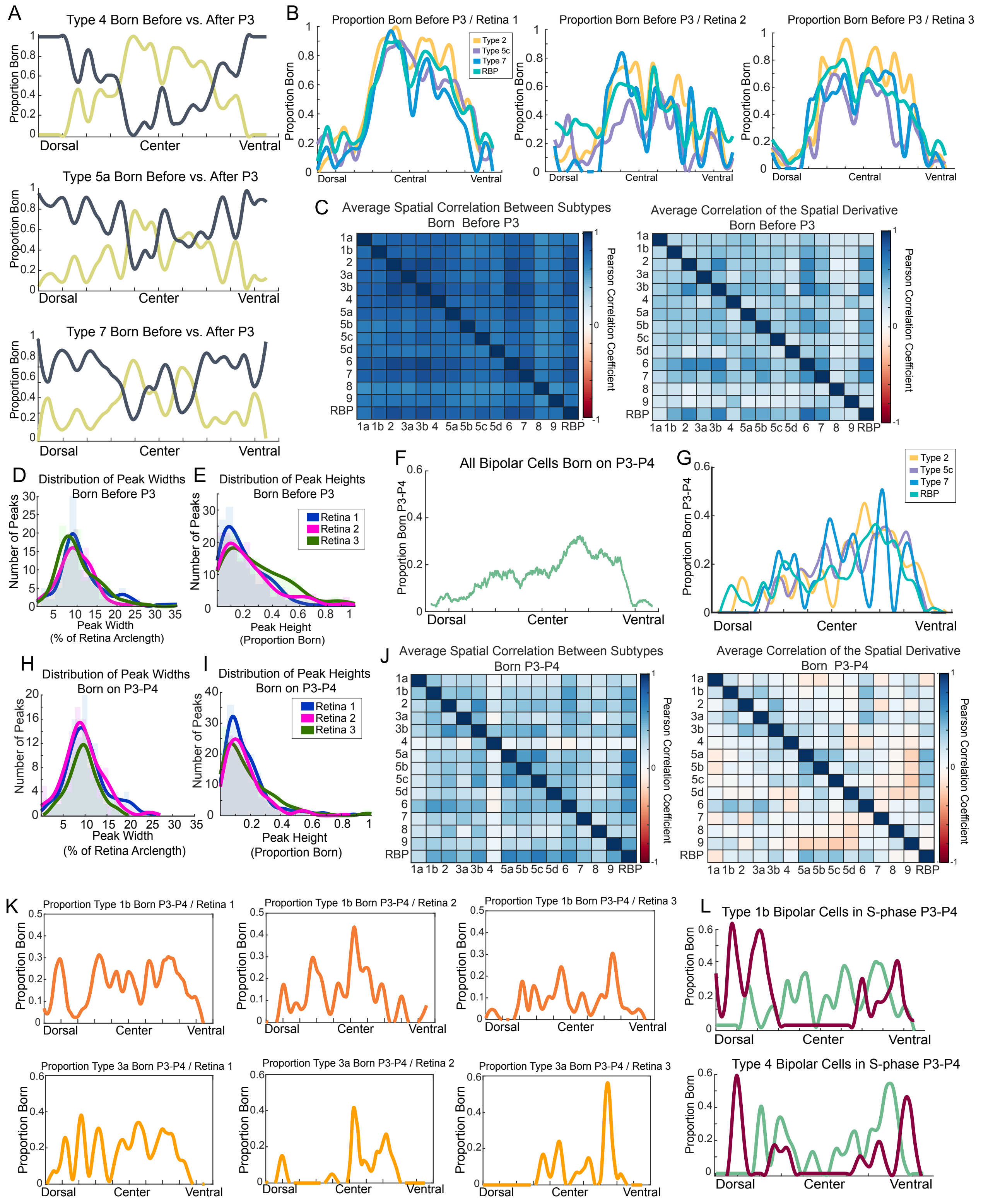
A
Type 4 Born Before vs. After P3
Type 5a Born Before vs. After P3
Type 7 Born Before vs. After P3
Proportion Born
Dorsal
Center
Ventral
B
Proportion Born Before P3 / Retina 1
Proportion Born Before P3 / Retina 2
Proportion Born Before P3 / Retina 3
Type 2
Type 5c
Type 7
RBP
Central
C
Average Spatial Correlation Between Subtypes Born Before P3
Average Correlation of the Spatial Derivative Born Before P3
Pearson Correlation Coefficient
1a 1b 2 3a 3b 4 5a 5b 5c 5d 6 7 8 9 RBP
D
Distribution of Peak Widths Born Before P3
Number of Peaks
Peak Width (% of Retina Arclength)
E
Distribution of Peak Heights Born Before P3
Retina 1
Retina 2
Retina 3
Peak Height (Proportion Born)
F
All Bipolar Cells Born on P3-P4
Proportion Born P3-P4
G
H
Distribution of Peak Widths Born on P3-P4
I
Distribution of Peak Heights Born on P3-P4
J
Average Spatial Correlation Between Subtypes Born P3-P4
Average Correlation of the Spatial Derivative Born P3-P4
K
Proportion Type 1b Born P3-P4 / Retina 1
Proportion Type 1b Born P3-P4 / Retina 2
Proportion Type 1b Born P3-P4 / Retina 3
Proportion Type 3a Born P3-P4 / Retina 1
Proportion Type 3a Born P3-P4 / Retina 2
Proportion Type 3a Born P3-P4 / Retina 3
L
Type 1b Bipolar Cells in S-phase P3-P4
Type 4 Bipolar Cells in S-phase P3-P4

**Figure S3. Automated classification of all bipolar subtypes *in situ*.** (**A**) Plot of Loess-smoothed SWAs of the local proportion of Type 4 (top), Type 5a (center), and Type 7 (bottom) born before (yellow line) vs. after P3 (blue line) in a single continuous section of Retina 1. (**B**) Overlay of the Loess-smoothed SWAs of the local proportion of Type 2, 5c, 7, and RBP born before P3 (BrdU-/EdU-) across 3 retina replicates, averaged across 3 sequential sections each. (**C**) Pair-wise Pearson correlations (averaged across 9 sections from 3 retinas) of the SWA of the proportion of each subtype born before P3 (curves shown in **(B)**) (left). To compare the local rate of change in birthdate patterns, the pair-wise Pearson correlations (averaged across 9 sections from 3 retinas) of the spatial derivatives of the SWA born before P3 were computed (derivatives of curves shown in **(B)**) (right). (**D**) Regions enriched for bipolar cell birthdates before P3 were considered "peaks", and were identified by a peak calling algorithm on the curves in (**B**). The distribution of peak widths (measured as % arclength along the retina space) is plotted across all subtypes and retinas. The distribution of peak widths (measured from trough to adjacent trough) are shown for three retina replicates. (**E**) The distribution of peak heights before P3 (measured as [[peak height] - [nearest trough height]]) is plotted across all subtypes and retinas. The distribution of peak heights are shown for three retina replicates. (**F**) SWA of the local proportion of all bipolars born on P3-P4 ($BrdU^{+}/EdU^{-}$ bipolar cells). (**G**) Overlay of Loess-smoothed SWAs of the local proportion of Type 4 (top), Type 5a (center), and Type 7 (bottom) born on P3-P4 in a single continuous section of Retina 1. (**H**) Regions enriched for bipolar cell birthdates on P3-P4 were considered "peaks", and were identified by a peak calling algorithm on the curves in (**G**). The distribution of peak widths (measured as % arclength along the retina space) is plotted across all subtypes and retinas. The distribution of peak widths (measured from trough to adjacent trough) are shown for three retina replicates. (**I**) The distribution of peak heights in the P3-P4 born population (measured as [[peak height] - [nearest trough height]]) is plotted across all subtypes and retinas. The distribution of peak heights are shown for three retina replicates. (**J**) Pair-wise Pearson correlations (averaged across 9 sections from 3 retinas) of the SWA of the proportion of each subtype born on P3-P4 (curves shown in **(G)**) (left). To compare the local rate of change in birthdate patterns, the pair-wise Pearson correlations (averaged across 9 sections from 3 retinas) of the spatial derivatives of the SWA born on P3-P4 were computed (derivatives of curves shown in **(G)**) (right). **(K)**) Examples of Loess-smoothed SWA curves of the proportion of Types 1b and 3a born on P3-P4 across 3 retina replicates. Curves were averaged across 3 sequential sections for each retina. **(L)**) Loess-smoothed SWAs of the proportion of Types 1b and 4 that were in terminal S-phase (green) or non-terminal S-phase (red) at the time of BrdU injection on P3.

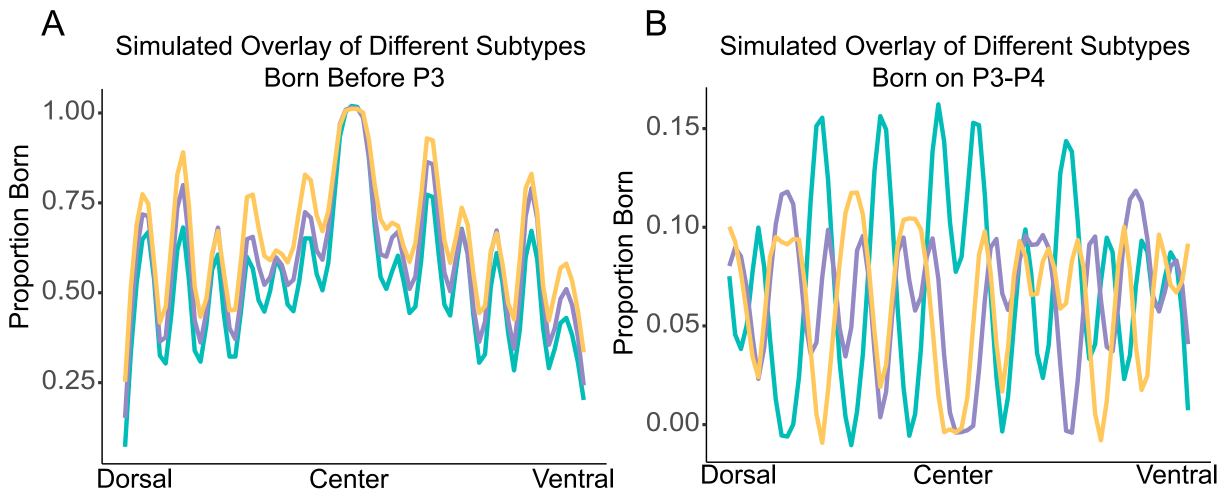


**Figure S4. Simulated birthdate patterns of bipolar subtypes based on a hierarchical genesis model**. (**A**) Bipolar subtype birthdates were simulated based on the hierarchical model described in Figure 7. Simulation results of SWAs of the proportion born before P3 for three different simulated bipolar subtypes. Analagous to real data presented in Supplementary Fig. S3B. (**B**) Simulation results of SWAs of the proportion born on P3-P4 for the same three simulated bipolar subtypes. Analagous to real data presented in Supplementary Fig. S3G.
